# Probing the modulation of enzyme kinetics by multi-temperature, time-resolved serial crystallography

**DOI:** 10.1101/2021.11.07.467596

**Authors:** Eike C. Schulz, Andreas Prester, David von Stetten, Gargi Gore, Caitlin E. Hatton, Kim Bartels, Jan-Philipp Leimkohl, Hendrik Schikora, Helen M. Ginn, Friedjof Tellkamp, Pedram Mehrabi

**Affiliations:** University Medical Center Hamburg-Eppendorf (UKE), Hamburg, Germany; Max-Planck-Institute for the Structure and Dynamics of Matter, Hamburg, Germany; Institute for Nanostructure and Solid State Physics, University of Hamburg, Hamburg, Germany; European Molecular Biology Laboratory (EMBL), Hamburg, Germany; Deutsches Elektronen Synchrotron (DESY), Hamburg, Germany

**Author notes:** These authors contributed equally to this work (Eike C. Schulz, Friedjof Tellkamp, Pedram Mehrabi).

**Keywords:** serial crystallography, 5D-SSX, multi-temperature crystallography, time-resolved crystallography, multi-dimensional crystallography

## Abstract

We present an environmental enclosure for fixed-target serial crystallography, enabling X-ray diffraction experiments in a temperature window from below 10 °C to above 70 °C - a universal parameter of protein function. Via 5D-SSX time-resolved experiments can now be carried out at physiological temperatures, providing fundamentally new insights into protein function. We show temperature-dependent modulation of turnover kinetics for the mesophilic *β*-lactamase CTX-M-14 and for the hyperthermophilic enzyme xylose isomerase.

## Introduction

Life has evolved to occupy a wide range of different environmental niches, ranging from glaciers and arctic deserts to the hot ponds at Yellowstone [1–3]. Although enzymes operate over a wide range of temperatures, the vast majority of protein structures have been acquired at cryogenic temperatures, to reduce the rate of radiation damage [4]. However, flash-cooling may introduce structural artefacts, which are absent at physiological temperatures, that may affect biological interpretation [5–8]. In addition, as proteins do not obtain a single unique conformation, but rather fluctuate around an energetic ground-state, the conformational dynamics of proteins are intimately linked to temperature, which is directly connected to their catalytic efficiency [9–12].The mismatch between data collection temperature and enzymatic activity optimum temperature becomes an acute problem during time-resolved structural studies that seek to provide detailed insight into catalytic mechanisms and allosteric regulation. While time-resolved structural studies are recently undergoing a resurgence, the majority of such experiments are carried out at ambient temperatures, regardless of the actual physiological or optimal temperature of the protein under study [13, 14]. A notable exception is the pioneering work on photoactive yellow (PYP) protein, where single-crystal time-resolved data were collected from -40 °C to +70 °C [15, 16]. Although it is a key environmental variable for mechanistic time-resolved analyses, considerable technological difficulties have prevented routine use of physiologically relevant temperatures during serial femtosecond (SFX) and serial synchrotron crystallography (SSX) experiments, until now. To address this challenge we have developed a modular environmental control system for serial crystallography, that enables precise control of the relative humidity to within a percent and temperature from below 10 °C and over 70 °C (**Fig.1 a, extended data**). This temperature window is large enough to encompass physiologically relevant temperatures of both mesophile and hyperthermophile enzymes. With full compatibility with our previously described liquid application method for time-resolved analyses (LAMA), this system permits multi-temperature time-resolved serial crystallography experiments [17, 18]. These so-called 5D-crystallography experiments [15] now enable a unique view into a hitherto difficult to structurally access realm of protein function, that directly connects out-of-equilibrium structures to energetic considerations. To demonstrate the applicability of our new environmental control system to a variety of enzymes, we have used a mesophilic enzyme, the extended spectrum *β*-lactamase (CTX-M-14) [19]) and a hyperthermophilic enzyme, xylose isomerase (XI) [20] as model systems. For both systems we show that a modification of the environmental temperature modulates the underlying enzyme kinetics, enabling resolution of alternate structural intermediates not observed at high occupancy at ambient temperatures.

**Fig. 1.**
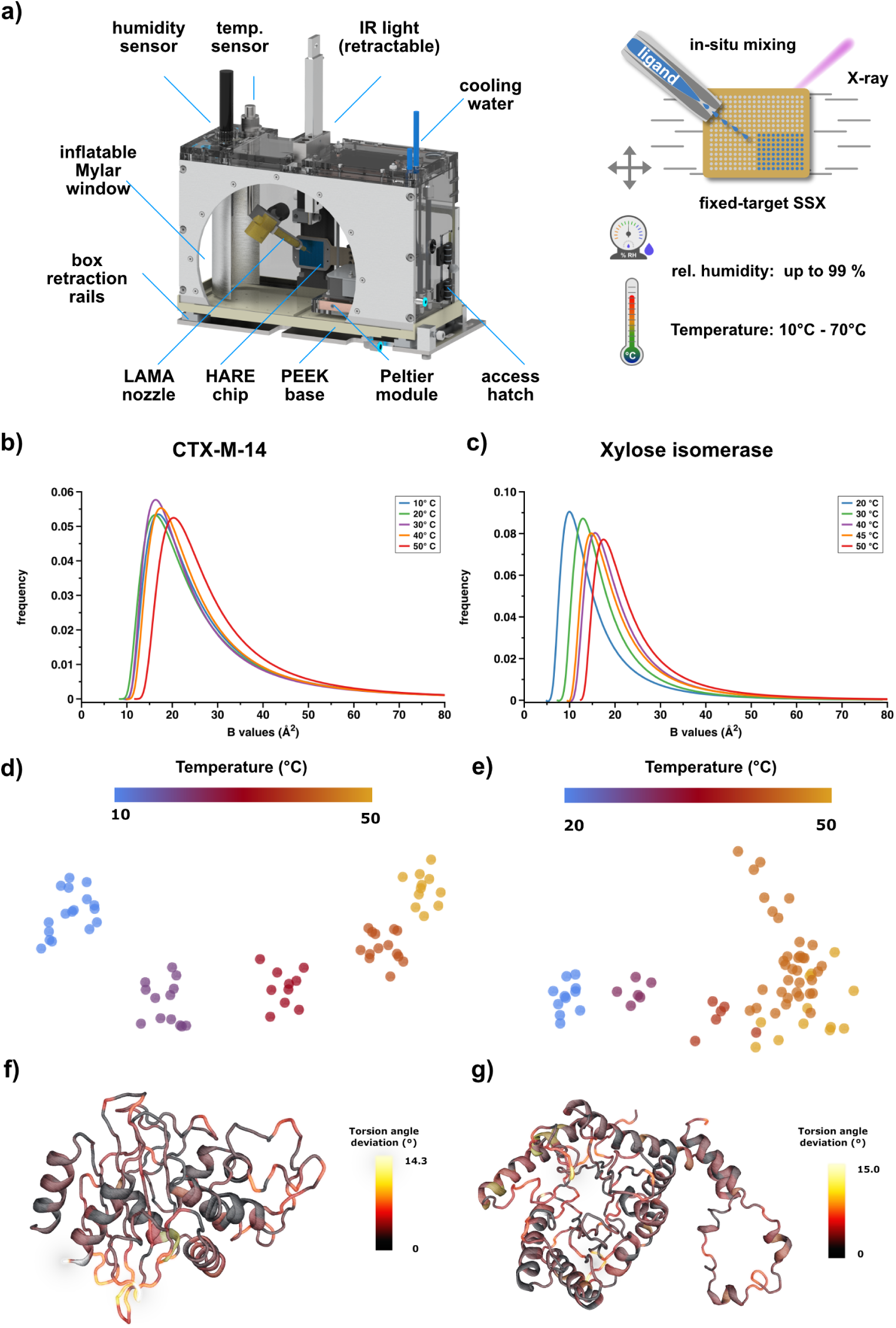
The environmental control box enables recording multi-temperature serial crystallography data. a) The environmental control box, the portal translation for the LAMA nozzle is hidden for clarity (further details in the extended data section). b,c) ADPs of CTX-M-14 and XI for models derived from data recorded at different temperatures, fitted to a shifted inverse gamma distribution. d,e) Atomic coordinate-based RoPE space for both proteins at each temperature [21]. Each point corresponds to an experimentally determined structure based on an image subset, showing the first two principal components. f,g) shows magnitude in torsion angle deviation corresponding to the first principal component of d) and e) respectively, plotted as a heatmap onto the backbone of the structures, illustrating the local structural response to temperature changes. In CTX-M-14 these are residues 52-54, 193-196, 226-231 and 251-255, while in XI the region around 25-26, 126-129 and 206-210 shows the largest torsion angle deviation.

## Results and Discussion

To benchmark the environmental control and systematically address different temperatures, five structures of both model systems were determined between 10 °C and 50 °C (Ext. Tab. 5, 6) (extended data). To assess the global variability of atomic positions in a comparable way, we determined the atomic displacement parameters (ADP), and fitted these to a shifted inverse gamma distribution as a function of temperature, as described previously [22]. For CTX-M-14 a shift in the ADP distribution can only be determined after a threshold temperature (ca. 50 °C), corresponding to its in-solution melting temperature and pairwise C*_α_*-RMSD (**Fig.1 b**, Ext. Fig. 6, 7). XI on the other hand has a much higher in-solution melting temperature (Ext. Fig. 6) and shows a monotonous ADP increase with higher temperatures (**Fig.1 c**). We hypothesise that the respective response in conformational variability reflects the adaptation to a mesophilic or a thermophilic temperature optimum, respectively. In other words, the higher flexibility observed for XI serves to accommodate the higher degree of molecular motions concomitant with higher temperatures, without compromising its functionality [20, 23]. However, ADPs do not only display local structural dynamics but also contain other sources of hierarchical disorder, such as lattice and molecular displacements [24], and may thus disguise subtle conformational changes. As changes in torsion angles are separated by lower energy barriers than changes in coordinates, they are more sensitive to conformational dynamics. Previously it was established that torsion angle deviations preserve temperature induced dynamics [21]. Therefore, we analysed the torsion angle distribution in both enzymes, to further investigate the response to altered environmental temperatures and delineate local structural changes. Clearly, both the CTX-M-14 as well as the XI sub-datasets assemble into temperature-specific clusters, emphasizing a coherent conformational space that corresponds to environmental temperature (**Fig.1 d,e**). However, analysis of the torsion angle dynamics also permits identification of local structural elements that directly respond to temperature changes. Thus, mapping the torsion angle deviation onto the backbone of the protein structures highlights hinge regions with particular flexibility (**Fig.1 f,g**). Collectively, these data unambiguously show a direct response of the conformational dynamics to the temperature inside of the environmental control box, and that these temperature changes can be effectively projected to structural changes.

Next, we analysed the effect of temperature on the catalytic activity of our two model enzymes. To assess thermal effects on catalysis, we equilibrated the crystals at a given temperature and triggered their reaction by adding the substrate solution *via* the LAMA-method [17] and monitored turnover at a constant time-delay of 3 s (CTXM-14) and 60 s (XI) after reaction initiation, respectively (extended data table 7 and 8). To minimize interpretation bias, we assembled the previously known, stable reaction intermediates into one structure and relied on constrained group refinement to determine the occupancy of the different overlapping sub-states at the respective delay time points (extended data) [25, 26]. Since different catalytic sub-states of CTX-M-14 can clearly be distinguished (unbound-state, acyl-enzyme intermediate, hydrolysed piperacillin product; **Fig. 2 a-c**), these data unambiguously show that both ligand diffusion as well as turnover kinetics of mesophilic enzymes can be modulated by temperature variation. While the piperacillin hydrolysis by CTX-M-14 is irreversible and progressively proceeds towards a product-bound state, glucose to fructose conversion by XI can also proceed in the backward direction [19, 23]. Accordingly, the system obtains an equilibrium between glucose and fructose over time. Snapshots along the reaction coordinate pathway would therefore reflect fractional occupancies of both species, mixed with open-chain reaction intermediates. As XI has an activity optimum at ca. 80 °C [20, 23], an increase of the temperature for a given delay time should shift the population towards the product side. In line with this hypothesis, the XI snapshots 60 s after reaction initiation show progressively decreasing occupancy for the glucose substrate and increasing occupancy for the fructose product with increasing temperature (**Fig. 2 d-f**). This demonstrates that thermal modulation enables deriving even nuanced shifts in reaction equilibria and thus detailed mechanistic distinctions as a function of the systems energy. This suggests that addressing catalysis at different temperatures enables simultaneous correlation of mechanistic and thermodynamic aspects of protein function. Conducting similar experiments on other systems will allow a general understanding how proteins exchange energy with their environment and how this is related to conformational dynamics and turnover. That is, multi-dimensional analyses are the foundation to experimentally determining free-energy landscapes of proteins in action and thus their unique catalytic pathways [15].

**Fig. 2.**
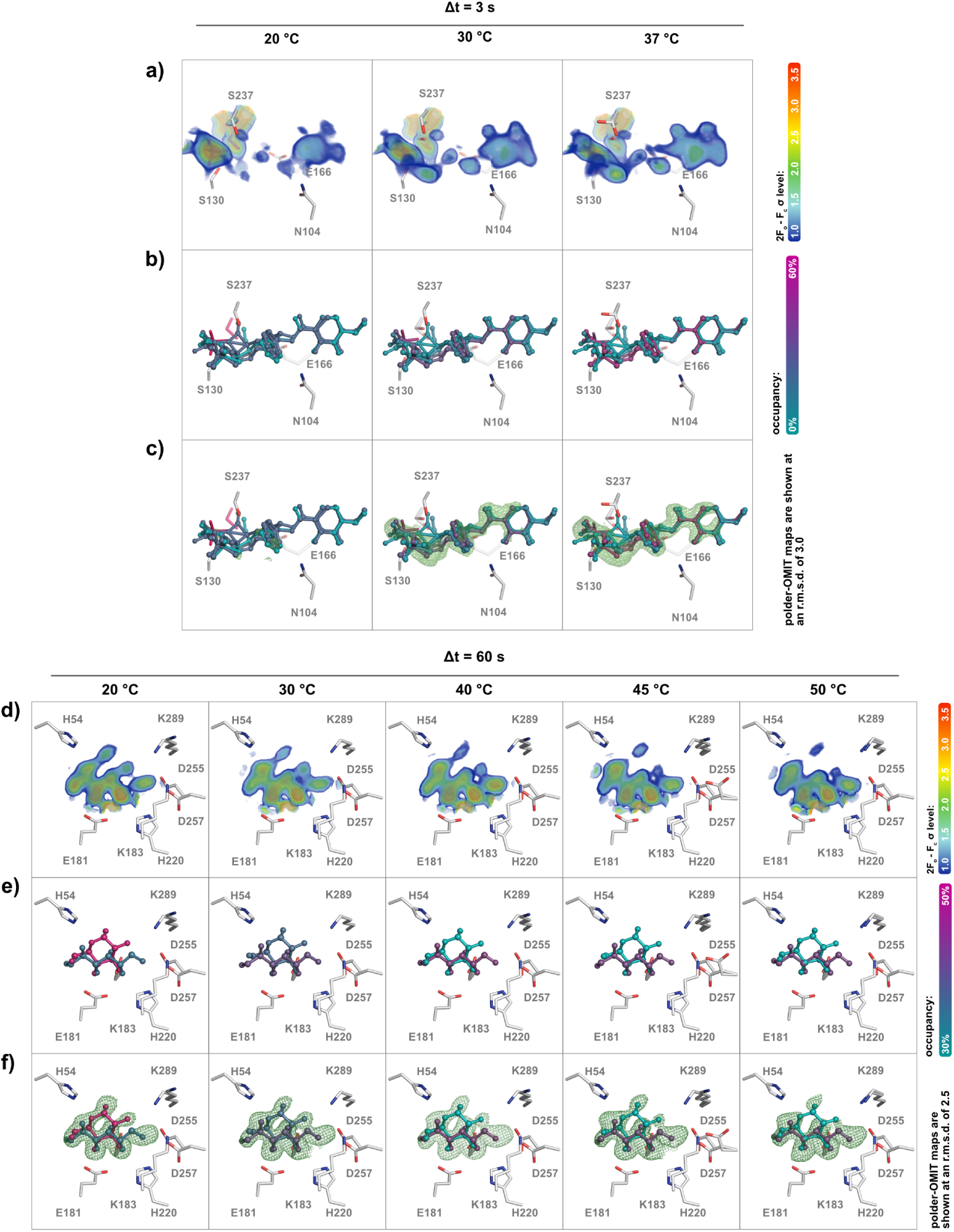
Altering enzyme kinetics by temperature modulation. a-c) CTX-M-14 active site, 3 s after reaction initiation at 20 °C, 30 °C, and 37 °C; d-f) XI active site, 60 s after reaction initiation at 20 °C, 30 °C, 40 °C, 45 °C, and 50 °C; a,d) 2*Fo − Fc* density shown at RMSD levels from 1.0 - 3.5, strongest density is shown in red; b,e) refined fractional occupancy level; c,f) polder-OMIT map shown around the ligands as a green mesh at the indicated RMSD levels.

In summary, our environmental control box permits charting new experimental space for the large number of enzymes that are amenable to time-resolved serial crystallography. *Via* temperature variation it enables the modulation of enzyme kinetics, thereby altering enzyme turnover and allowing for a more in-depth characterization of enzymatic mechanisms. We anticipate that such experiments will contribute to the changing role of structural biology to enable a comprehensive understanding of conformational dynamics and its role in protein function in the future.

## Material and Methods

### Environmental control system

For a detailed description of the environmental control box design and characterisation of the temperature and humidity controls, please refer to the extended data section. Briefly: the environmental control box consists of a rail-mounted, retractable, perspex housing with cycloolefin copolymer (COC) and mylar X-ray inlet and outlet windows, respectively. Constant relative humidity and temperature can be achieved by two independent closed loop control circuits, whereby the former is controlled by a flow of humidified air from a hot water bath. Two interchangeable modules are used to control the temperature. Module-1 contains water-cooled Peltier elements, while module-2 consists of simple heating resistors. The modules cover the ranges from +7 °C to +50 °C and +50 °C to +70 °C, respectively. All information that is required for reproduction of the hardware, including 3D-PDFs, CAD files, step-files, electronics, and a bill of materials can be obtained under following DOI: 10.5281/zenodo.12758835

### Sample preparation

Xylose isomerase was purified as described in detail in the extended data section. For crystallisation, the protein was then concentrated to 80 mg/ml using a 10 MWCO concentrator (Sartorius). Subsequently, microcrystals were obtained by vacuum induced crystallization, as described previously by Martin et al. [27], in XI crystallization buffer (35% (w/v) PEG 3350, 200 mM LiSO_4_ and 10 mM Hepes/NaOH, pH 7.5). Sufficient microcrystal for a typical HARE chip were prepared from 25 µl protein solution (80 mg/ml for xylose isomerase) combined with 25 µl crystallization buffer. For droplet injection 1 M glucose solution was prepared in ddH_2_O prior to data collection and stored at room temperature.

CTX-M-14 was purified as described previously [28]. The CTX-M-14 solution (22 mg/ml) was mixed with 45% (v/v) crystallising agent (40% (w/v) PEG 8000, 200 mM LiSO_4_, 100 mM sodium acetate, pH 4.5) and with 5% (v/v) undiluted seed stock solution to induce micro-crystallization. This resulted in crystals with a homogeneous size distribution of 11-15 µm after approximately 90 minutes. To stop further crystal growth crystals were centrifuged at 200 x g for 5 min and the supernatant was replaced with a stabilisation buffer (28% (w/v) PEG 8000, 140 mM LiSO_4_, 70 mM sodium acetate, 6 mM MES pH 4.5, 15 mM NaCl).

### Determination of the melting temperatures

The melting temperatures of CTX-M-14 and XI were determined in four different buffer systems each. The buffers tested were the protein storage buffer (buffer 1), crystallization buffer (buffer 2), crystallization buffer without PEG (buffer 3) and activity assay buffer (buffer 4). The proteins were diluted to 1 mg/mL in the respective nanoDSF buffers and incubated for 20 min on ice. Standard grade nanoDSF capillaries (Nanotemper) were loaded into a Prometheus Panta (Nanotemper) controlled by PR. PantaControl (x64). Excitation power was adjusted to 25% and samples were heated from 20 °C to 95 °C with a slope of 1 °C/min. All samples were examined in triplicates and error bars represent standard deviations. Buffer details are listed in the extended data section.

### Experimental setup and data collection

Diffraction data were collected at the EMBL endstation P14.2 (T-REXX) at the PETRA-III synchrotron (DESY, Hamburg) with an X-ray beam of 10 × 7 µm (H×V) on an Eiger 4M detector (Dectris, Baden-Daettwil, Switzerland). Data collection was conducted as previously described [18] within the environmental control box mounted on the T-REXX endstation. Briefly: microcrystals mounted in a HARE-chip solid target containing 20,736 wells were moved through the X-ray beam using a 3-axis piezo translation stage setup (SmarAct, Oldenburg, Germany) [29]. Time delays for time-resolved data collection were generated *via* the HARE method and reaction initiation was achieved *via in-situ* droplet injection via the LAMA method, as previously described in detail [17, 18, 29]. As a broad guideline approximately 5000 still diffraction patterns were recorded per time-point as previously determined to be sufficient [30].

### Data processing and structure determination

Diffraction data were processed using the CrystFEL v0.10.0 package [31]. Structures were solved by molecular replacement using PHASER with our previously determined XI and CTX-M-14 structures as a search model with one molecule in the asymmetric unit (PDB-ID: 6RNF, 6GTH) [32]. Structures were refined by iterative cycles in *phenix.refine* and manual model building of additional and disordered residues in COOT-v0.8 [33–35]. Occupancy refinement was carried out in *phenix.refine* by defining constrained occupancy groups of ligand-free and ligand-bound proteins, and refining against all states simultaneously. Further details are given in the extended data section.

### Electron density figures

POLDER-OMIT maps were generated using *phenix.polder*, omitting the ligand residues in the constrained occupancy groups [36]. Molecular images were generated in PyMOL [37]. Further details are given in the extended data section.

### Fitting ADPs to a shifted inverse gamma distribution

The isotropic ADP frequency histograms of each structure were fitted with a three parameter Shifted Inverse Gamma Distribution (SIGD) function as shown previously by Masmaliyeva et al [22]. However, in contrast to the maximum-likelihood estimation using the Fisher scoring method applied previously [22], parameter estimation and optimization was carried out by using non-linear least squares with the python module scipy.optimize.curve_fit. To produce reasonable estimates, necessary parameter restraints [22], were applied during the estimation process. Briefly, the starting value of the shift parameter was taken to be equal to 90% of the minimum of the ADPs in the PDB file.

### Sub-dataset splitting and analysis with RoPE

For the RoPE analysis, each dataset was split into subsets of at least 2000 diffraction patterns each using *partialator* [31], which were then independently refined with DIMPLE [38], without human intervention. This allows us to assess the relative contributions of random fluctuation and genuinely temperature-dependent changes within the crystal structure, as employed previously [39]. The output of these PDBs was analysed in RoPE [21] to show the first two principal components of atomic coordinate differences, and corresponding temperature metadata.

## Acknowledgments

All multi-temperature SSX data were collected at endstation P14.2 (T-REXX) operated by EMBL Hamburg at the PETRA-III storage ring (DESY, Hamburg, Germany). We would like to thank our colleagues G. Bourenkov, M. Agthe, and A.R. Pearson for their continuous support, helpful discussions and critical reading of the manuscript.

## Author Contributions

P.M. and E.C.S. designed the experiments; E.C.S. and P.M. performed the experiments with support from D.v.S. and F.T.; P.M., A.P., C.E.H. and K.B. prepared protein and the protein crystals; E.C.S., P.M., J.P.L. and F.T. designed and built the environmental control box; D.v.S, A.P, E.C.S. and P.M. processed and analyzed the diffraction data; P.M., E.C.S., H.S., J.P.L. and F.T. characterized the control box; P.M. and E.C.S. wrote the manuscript; All authors discussed and corrected the manuscript.

## Declarations

Competing Financial Interests Statement: The authors declare no competing financial interests.

## Extended Data

### Extended methods

#### Protein purification

Xylose isomerase (Uniprot ID: P24300) was cloned into pET-24a(+). The construct was transformed into *E. coli* strain BL21 (DE3) Gold and grown in TB medium supplemented with 25 µg/mL kanamycin at 37 °C until an OD_600_ of 1.0 - 1.2 was reached. Protein expression was induced by addition of 1 mM IPTG, and the cells were incubated further at 18 °C for 16 h. The cells were harvested by centrifugation (7000 g, 15 min, 4 °C) and resuspended in lysis buffer (50 mM HEPES pH 7.5, 500 mM NaCl, 5% (v/v) glycerol, 5 mM imidazole, 5 units/mL DnaseI, and protease inhibitor). After the cells were lysed by sonication, undisrupted cells and debris was separated by centrifugation (30,000 g, 1 h, 4 °C). The supernatant was applied to a 5 mL HisTrap FF (Cytiva), washed with wash buffer (50 mM HEPES pH 7.5, 500 mM NaCl, 5% (v/v) glycerol, 30 mM imidazole), and XI was eluted in elution buffer (50 mM HEPES pH 7.5, 500 mM NaCl, 5% (v/v) glycerol, 250 mM imidazole). Protein containing fractions were pooled and HRV-3C protease was added to the eluate (0.3 mg HRV-3C protease for material derived from a 1 L culture). The sample was dialysed overnight against SEC buffer (50 mM HEPES pH 7.5, 150 mM NaCl) at 4 °C. Negative IMAC was performed to recover the cleaved XI. The protein was concentrated to 3 mL using a 10 MWCO concentrator (Sartorius) and applied to a Superdex 200 HiLoad 16/60 column (Cytiva) equilibrated with SEC buffer. Fractions containing the protein were pooled and a buffer exchange into crystallisation buffer (10 mM HEPES pH 7.5) was performed using a PD10 column (Cytiva).

#### Group occupancy refinement settings

Following group occupancy refinement setting have been used for CTX-M-14:

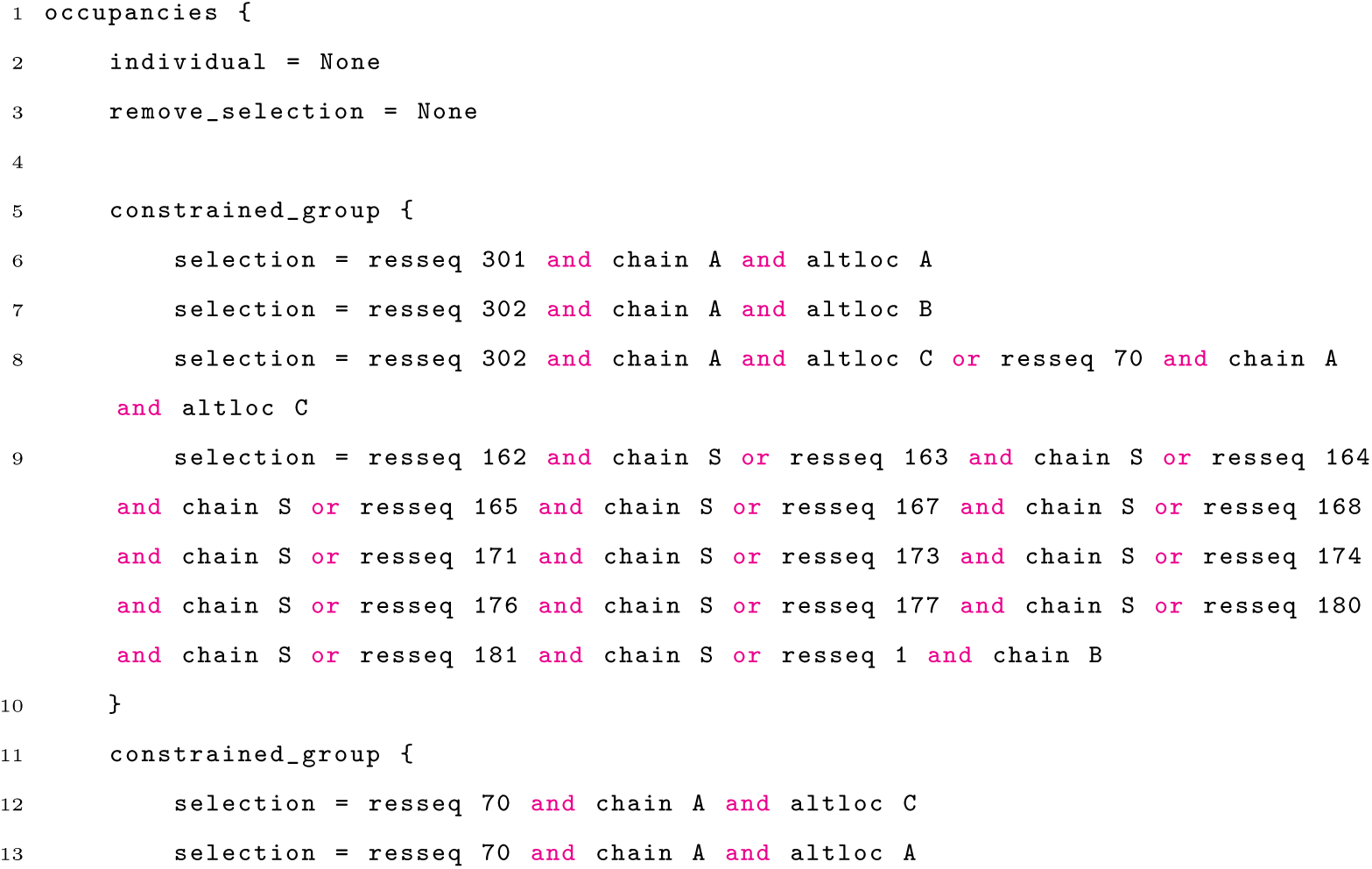
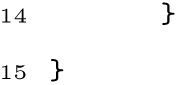

Following group occupancy refinement setting have been used for XI:

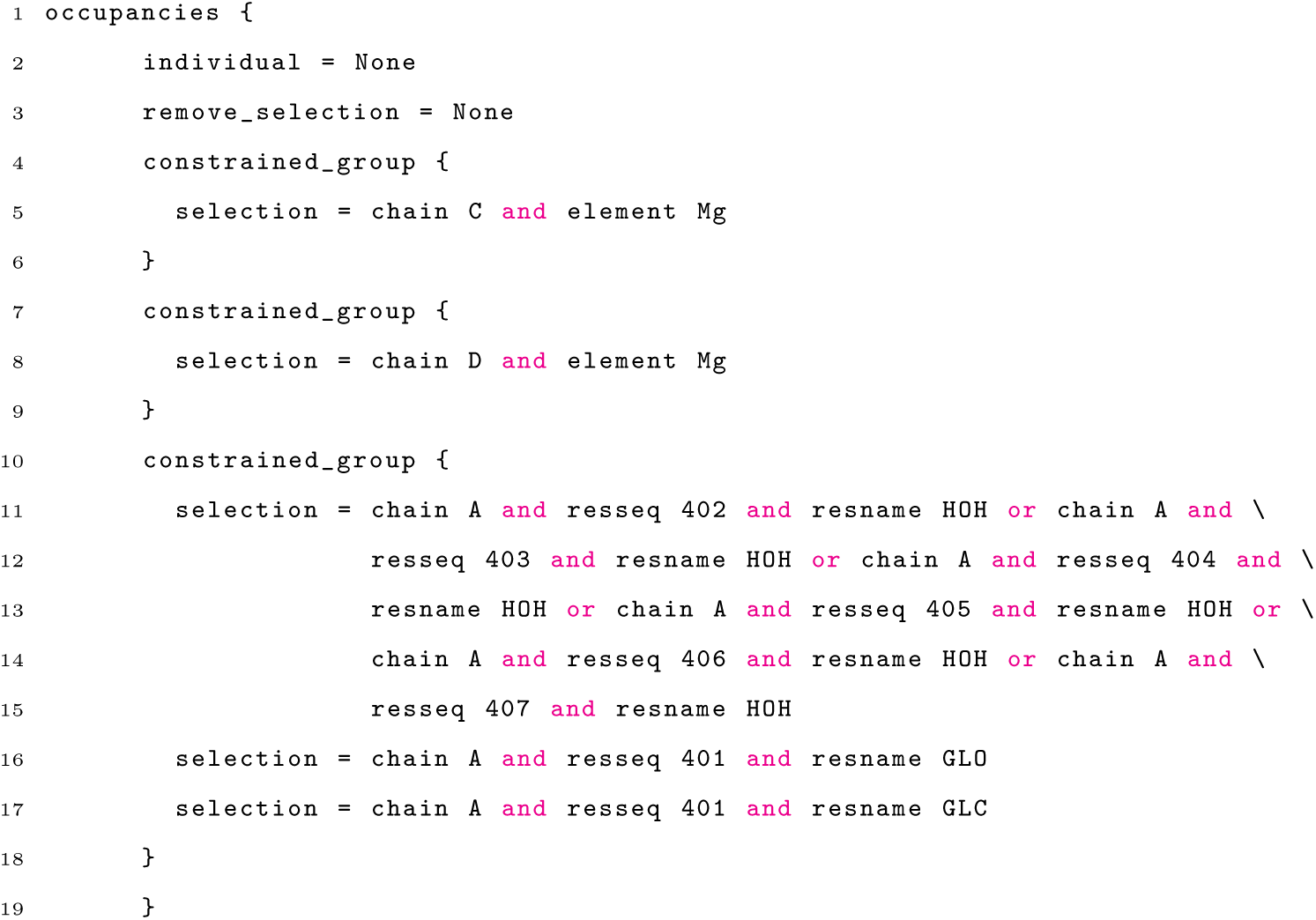

#### Electron density figure details

The 2*F_o_ − F_c_* electron density as volume elements with different RMSD levels were generated in PyMol *via* the volume command (`volume volumename, mapname, level, ligandname, carve=1.4`), where color and RMSD-level settings were according to Ext. Tab. 1 have been used. Ligand occupancy was displayed via following command: `spectrum q, teal_hotpink, imp, minimum=0.1, maximum=0.6` for CTX-M-14 and `spectrum q, teal_hotpink, imp, minimum=0.3, maximum=0.5`.

**Ext. Tab. 1.** PyMol color settings to display volume elements.

| Color | map RMSD. | R | G | B |
| --- | --- | --- | --- | --- |
| Blue | 1.0 | 19 | 43 | 157 |
| Cyan | 1.5 | 153 | 219 | 223 |
| Green | 2.0 | 101 | 198 | 66 |
| Yellow | 2.5 | 255 | 240 | 0 |
| Orange | 3.0 | 252 | 131 | 0 |
| Red | 3.5 | 245 | 38 | 0 |

### Environmental control box

#### Design

To maintain a controlled humidity environment for our hit-and-return (HARE) chip setup including the liquid-application-method for time-resolved crystallography (LAMA) [18, 30], required the development of a solution that could accommodate this experimental setup. To this end we have constructed a modular, compact environmental control box that encloses our previously described chip setup including the LAMA droplet injector nozzle on a footprint of 118 mm x 283 mm (Ext. Fig. 1, 2) [17]. Humidity control is achieved by flowing dry air either directly into the box, or first passing through a water bath (20-95 °C), with the proportion of gas through each channel controlled by a toggling ball valve in a (proportional-integral-derivative) PID-feedback loop. The water-bath is connected to the environmental control box via a silicone hose, to which a heating belt is attached preventing condensation. The set point can be achieved with high accuracy, enabling humidity control within 1 percent points of relative humidity, which allows to perform controlled crystal dehydration if that is required. Many beamlines are equipped with a temperature control solution via a gas stream that is directed at the sample, often encompassing wide temperature windows. Interestingly, however, a combined temperature-humidity control is rarely found. Such a situation mandates the use of e.g., glass capillaries to maintain the crystals in a humid environment, which complicates time-resolved applications that are based on in-situ mixing. Historically, flow-cells were developed for time-resolved applications that allowed for in-situ mixing experiments with single crystals and for trapping reaction intermediates on comparably slow time scales [40–42], which were recently extended to SSX experiments [43]. However, to the best of our knowledge these were not applied to multi-temperature experiments. Serial crystallography experiments can also be conducted in controlled, closed-boundary environments as demonstrated by the drop-on-demand device, which permits humidity control and fully anaerobic experiments via the exchange of the surrounding atmosphere [44]. In addition to humidity control, the temperature within our enclosure can be adjusted anywhere within the range of approximately 7 °C to above 70 °C. An aluminum air-stream reflector directs the stream of humid air around the chip. In order to enable effective control over this wide temperature range, the box uses two interchangeable modular temperature control units (Ext. Fig. 2). Module-1 covers a temperature range from approximately +7 °C to approximately +50 °C, while module-2 covers a temperature range from approximately +50 °C to over +70 °C. Rapid exchange of the modules is possible without tools, enabling switching between different temperature regimes within a few minutes. Module-1 are water-cooled Peltier-elements that enable active cooling or heating of the interior of the box. To ensure that the temperature is equilibrated across the box, the module is equipped with fans. Cooling water and electric power are fed in through the top side of the module. The heating element in module-2 is a power resistor network that disseminates sufficient heat to increase the interior temperature of the box to over 80 °C, while the relative humidity can be sustained at over 95%. Temperature control is achieved by a PID controller that sets the current through the Peltier elements and resistor network, respectively, to maintain the target temperature. The base plate of the box is made from durable polyether ether ketone (PEEK). To drain condensation water, several cotton wicks are fixed around the bottom corners and connected to an active pumping system that quickly drains excess liquid from the box. The sides and the lid of the box are made of 6 mm thick acrylic glass, while the rear panel (X-ray side) is made of acrylic glass and polyoxymethylene (POM). As an X-ray entrance window, an 8 mm opening in the POM is covered with two spaced layers of COC foil, about 1 mm apart. To enable easy replacement of the X-ray entrance window, the COC foil is placed on a magnetically mounted ring that tightly seals the inside of the box. While the right panel is solid, the left panel has a feedthrough for the humidity tube, and an access hatch through which HARE chips can be loaded onto the sample translation stage. The front panel (detector side) is an aluminum frame with a 190 mm opening. This is sealed with two X-ray transparent Mylar foils (6 µm), as an exit window for the diffracted beam. To reduce condensation on this window, the space between the Mylar foils is continuously flushed with warm air. The lid of the box contains feedthroughs for the humidity and temperature sensors, as well as for the electropneumatic retractable, infrared (IR) backlight, the LAMA-nozzle lever, and an access port for heating module exchange. The translation stage system implemented at the T-REXX endstation is not humidity resistant and therefore has to be kept outside of the box. Hence, the translation stages are connected to the box via a flexible bellow, custom cut from two layers of commercial plastic wrap. On the inside, the bellow is sealed between the translation stages and the kinematic mount for the chip holder [29, 45]. To reduce the heat capacity of the chip holder and thus achieve faster temperature equilibration, the previously described aluminum chip holder was redesigned from PEEK, providing the same functionality at a lighter weight [29]. The LAMA nozzle is attached *via* a kinematic mount to a retractable lever that enables retraction of the nozzle from its injection position during chip exchange. To avoid the unnecessary opening of the box, which might lead to temperature and humidity fluctuations, the nozzle retracts into a parking position under the lid of the box. Fine alignment of the LAMA nozzle in the injector position is achieved *via* motorized translation stages (Thorlabs). The whole box system is mounted on rails, residing on a stainless-steel baseplate that can be moved between “data collection” or “beam location” position, where the latter allows to use the X-ray scintillator built into the beam-shaping device (BSD; Arinax, Moirans, France). Operation of the serial crystallography environmental control enclosure is achieved *via* an external control unit where temperature and humidity parameters are electronically set (Ext. Fig. 3). Parameters needed to achieve certain conditions were calibrated and are listed in **Ext. Tab. 2**.

**Ext. Fig. 1.**
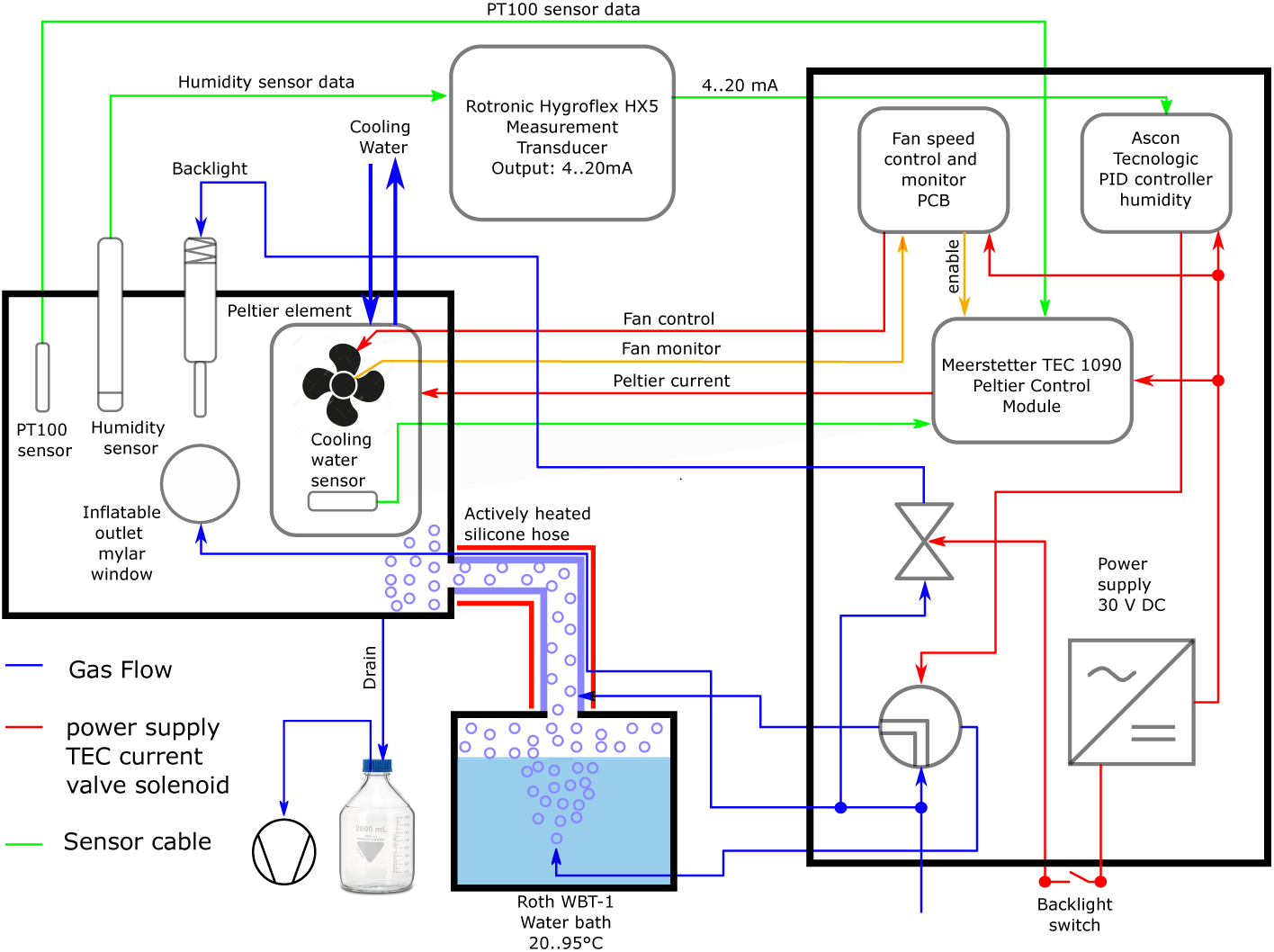
Schematic representation of the environmental control box setup. Gas flow is indicated by blue lines, power supply is red, sensor cables are depicted in green, and cooling water is displayed in pink.

#### Characterization of temperature and humidity stability

To characterize how the environmental parameters can be controlled within the box, we recorded temperature and humidity changes as a function of time in 30 s intervals. The temperature was modified as a step function, with the humidity target set to 95%.

**Ext. Fig. 2.**
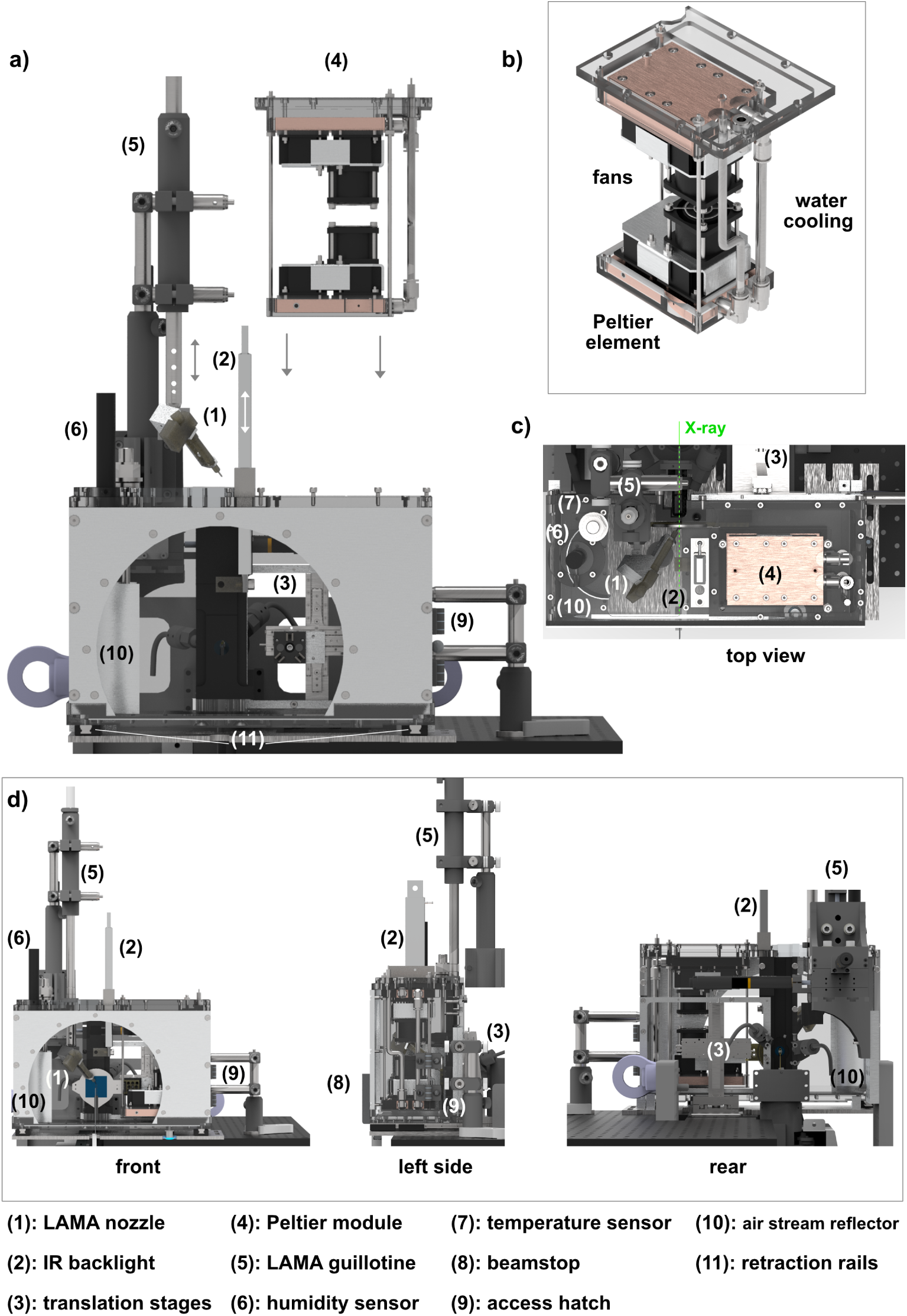
The Serial Crystallography Environmental Control Box. a) overview of the environmental control box, with retracted LAMA nozzle and Peltier module-1. b) closeup of the Peltier module-1, c) top-view providing an overview of the arrangement inside the box, d) front-, side- and rear-view of the box. Note: for clarity module-2 and some technical elements of the box (e.g. tubing, electric connections etc.) or the beamline are not shown or described in detail. Elements mentioned in the text are numbered and shown in the figure legend.

**Ext. Fig. 3.**
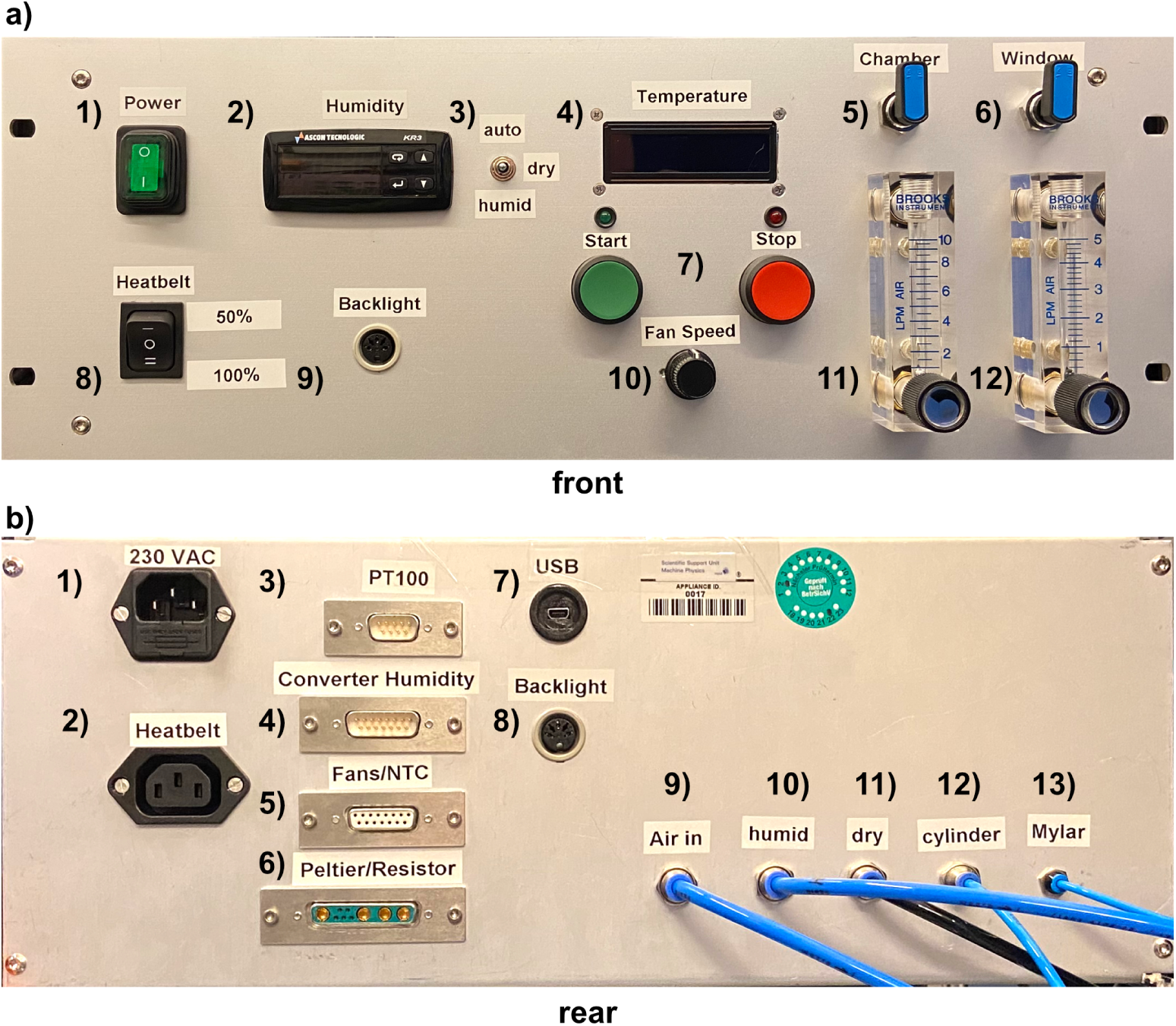
Control unit of the serial crystallography environmental control box. a) front panel: 1) main power switch; 2) humidity control/display; 3) air flow switch; 4) temperature display; 5) airflow on/off switch; 6) inflatable front-window flow on/off switch; 7) Temperature control start/stop button; 8) heat-belt switch; 9) socket for IR backlight switch; 10) fan-speed regulator; 11) air-flow valve; 12) front-window flow valve. b) rear panel: 1) main power inlet; 2) heat belt power outlet; 3) PT100 temperature sensor; 4) humidity sensor; 5) fan connection; 6) Peltier/resistor connection; 7) USB connection to PC; 8) backlight connection; 9) pressurized air in; 10) air flow to water bath; 11) dry air outlet; 12) backlight cylinder air outlet; 13) inflatable front-window air outlet.

While the humidity rapidly equilibrates throughout the box, temperature gradients closer to the walls needs to be avoided. In addition to the overall temperature inside the box, measured near the panel opposite to the access hatch, we also monitored the temperature at two additional locations, directly on the surface of the backside of the HARE chip and directly above the humidity sensor (Ext. Fig. 4). To assess how effectively different temperatures in the box can be achieved and maintained, we characterized the temperature increase from 7.5 °C to 80 °C. The data show that for both temperature control modules the humidity values quickly reach the target values. Over a temperature window of almost 70 °C the humidity remains stable within 2.5% of the set point. Analysis of the deviation of the chip temperature from the box temperature shows that the chip temperature follows the box temperature with a median difference of 0.7 °C, over a temperature window of almost 70 °C. Temperature and humidity typically equilibrate across the box and the chip within 10-15 minutes. We also examined the reliability of maintaining environmental set points during X-ray data-collection. To this end we collected X-ray diffraction data at 20 °C, 40 °C, 55 °C and 80 °C, for ca. 120 minutes and recorded temperature and humidity values in 30 s intervals during this period (Ext. Tab. 3). Remarkably, during the data collection the target humidity could be maintained within approximately 1%, while the temperature remained stable within approximately 0.5 °C. Clear deviations from this behavior are only observed during chip exchange, when the hatch of the box is opened and during the subsequent re-equilibration time while the environment stabilises. Re-equilibration of the environmental parameters could be achieved within 10-15 minutes, depending on the duration of the manual chip exchange. In conclusion, these data show that after an equilibration time of approximately 10-15 minutes the environment in the control box has reached its target value, and can be maintained throughout extended periods of time, well beyond the typical data collection time of a chip (ca. 30 minutes). This enables collection of serial X-ray diffraction data at a variety of different temperature and humidity levels with high accuracy and precision.

**Ext. Tab. 2.**
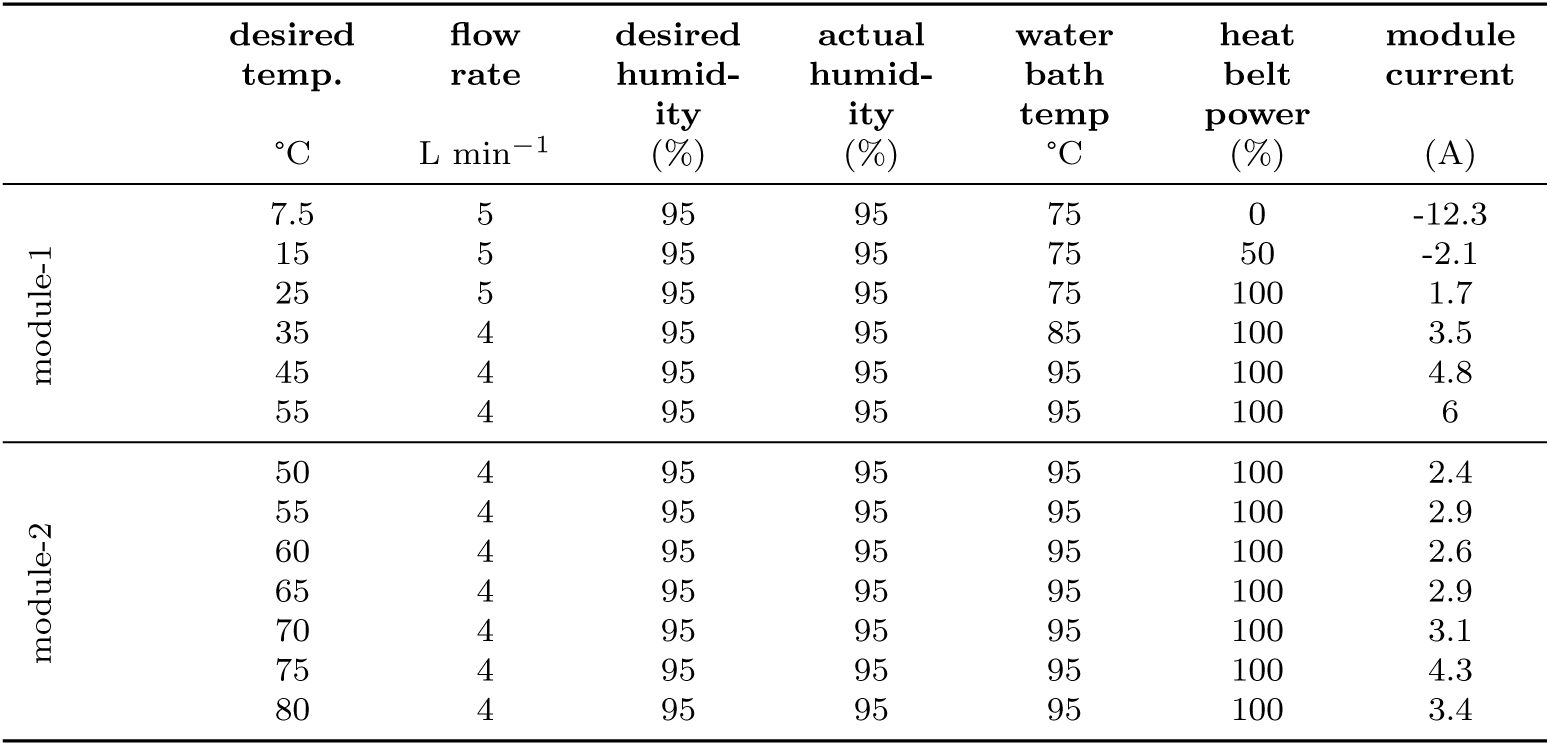
Environmental control parameter settings.

**Ext. Tab. 3.** Environmental parameters during X-ray diffraction data collection. *values are derived from a long-term, (10 h) data-collection

| nominal<br>temp. (° C) | nominal<br>rel.<br>humidity<br>(%) | measured<br>rel.<br>humidity<br>(%) | RMSD (%) | measured<br>temp. (° C) | RMSD (°<br>C) |
| --- | --- | --- | --- | --- | --- |
| 20 | 95 | 95.4 | 0.5 | 19.8 | 0.2 |
| 40 | 95 | 95.5 | 0.9 | 40.3 | 0.3 |
| 55 | 95 | 95.5 | 0.7 | 55.3 | 0.7 |
| 80 | 95 | 95.2 | 0.7 | 80.1 | 0.3 |
| 20* | 95* | 95.2* | 0.7* | 19.8* | 0.2* |

#### Humidity controlled SSX

A hallmark of protein crystals is their large solvent-content, which is typically in the range between 40% and 70% of the crystal volume. An advantage of this property is that proteins generally retain their function even in the crystalline state [46, 47]. A commonly known disadvantage is, however, their sensitivity to changes in environmental humidity, which in addition to the higher rates of radiation damage associated with higher temperatures makes routine data collection at even ambient temperatures a difficult task. Accordingly, starting with traditional wax-enclosed glass capillaries, environmental control has consistently been a key aspect of macromolecular crystallography, and several solutions have been developed to maintain crystal hydration for single crystals [4, 48]. Open- and closed boundary environmental control solutions have been developed since the advent of structural biology. In the simplest instances, closed boundary devices include glass capillaries, which contain protein crystals and typically a drop of mother liquor to sustain a humid atmosphere during data collection [48]. More modern variations of this classic solution are the many fixed-target serial crystallography environments, which protect protein microcrystals against evaporation by some form of X-ray transparent window material [49]. For single crystal experiments a variety of solutions, sometimes for advanced parameter control, such as humidity, temperature, and electric fields have also been developed over the years [50–52]. With the onset of cryo-crystallography, larger boxes were developed, which sometimes enclosed the stream of cryogenic gas in a dry atmosphere to prevent ice-formation during datacollection [53]. However, for single-crystal experiments the majority of environmental control solutions fall into the open-boundary category, such as placing the crystal in an vapor stream with controlled humidity [51, 54–58]. Historically, the water content of protein crystals was controlled by post-crystallization treatments via chemical dehydration prior to crystal freezing [59, 60]. However, controlling the humidity around the mounted crystals enables the convenient identification of the optimal conditions for a particular sample during an X-ray diffraction experiment [51, 54–58]. Adjusting the relative humidity either prior to or during data collection can improve several aspects of data quality (resolution, mosaicity and anisotropy) [54, 55, 57, 61–66]. To estimate the effect of the environmental humidity on diffraction data quality we monitored the unit-cell size of XI as a function of decreasing humidity. We started data collection at a relative humidity of 95% and reduced the humidity in steps of 5% per compartment row on the chip (**Fig. 5**). With decreasing humidity the crystals do not simply cease to diffract but undergo a change in unit-cell size. While at or above a relative humidity of 95% most diffraction patterns could be indexed with a larger unit-cell (a = 94.2 Å, b = 99.3 Å, c = 103.1 Å; *α, β, γ* = 90.0°), but the proportion rapidly changed to a smaller unit cell (a = 94.6 Å, b = 99.4 Å, c = 87.5 Å; *α, β, γ* = 90.0°) as the relative humidity dropped from 90% to 75%. If the humidity is reduced even further the micro-crystals cease to diffract, presumably due to complete dehydration.

**Ext. Fig. 4.**
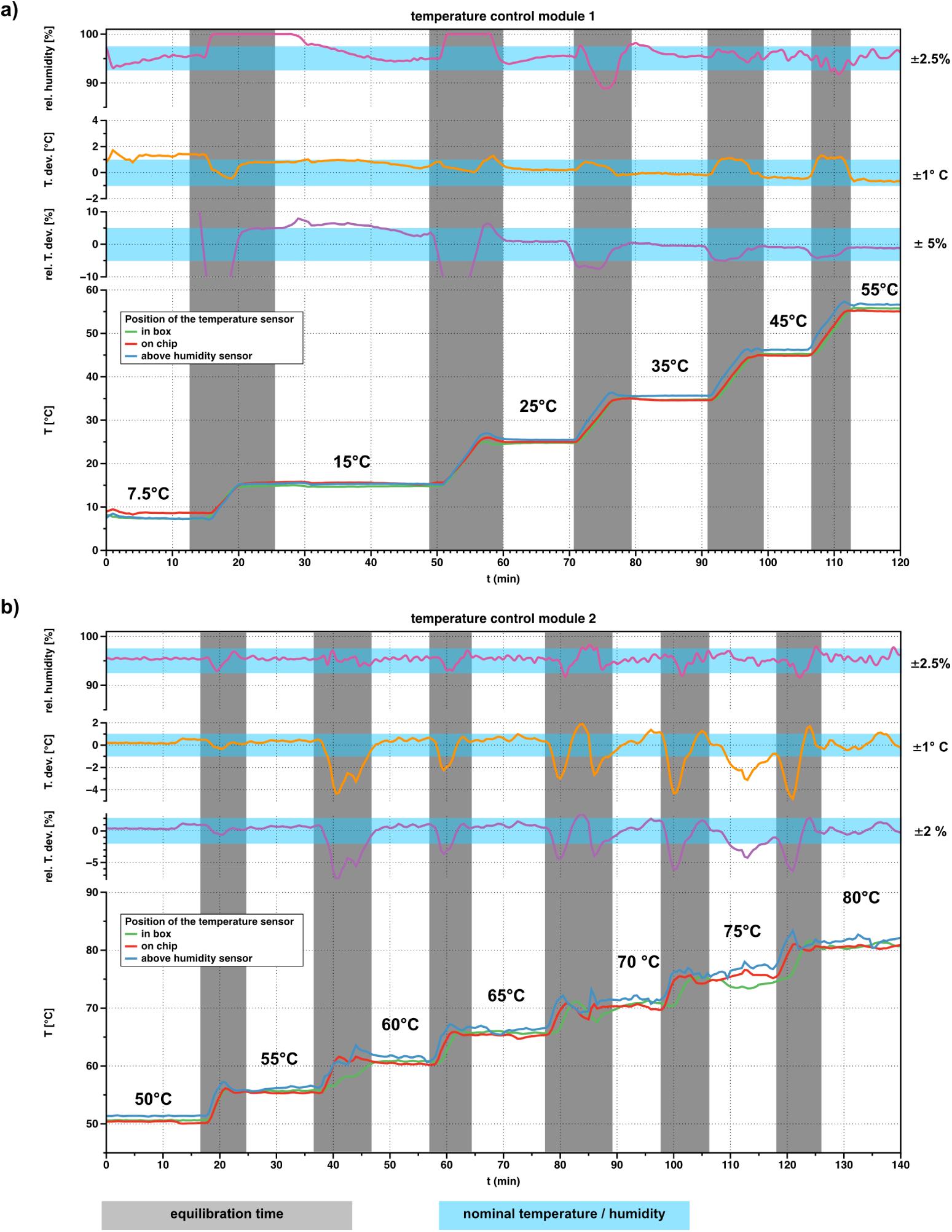
Characterization of the environmental control. a) module-1: 7.5 °C – 55 °C; b) module-2: 50-80 °C; The temperature was successively increased from 7.5 °C to 55 °C and 50 °C to 80 °C respectively. The temperature was measured at three positions inside of the chamber: green inside the box near the panel opposite to the access hatch, blue directly above the humidity sensor, red directly on the chip. The target humidity was set to 95%. The grey bars indicate the equilibration time, blue bars indicate the target window.

**Ext. Fig. 5.**
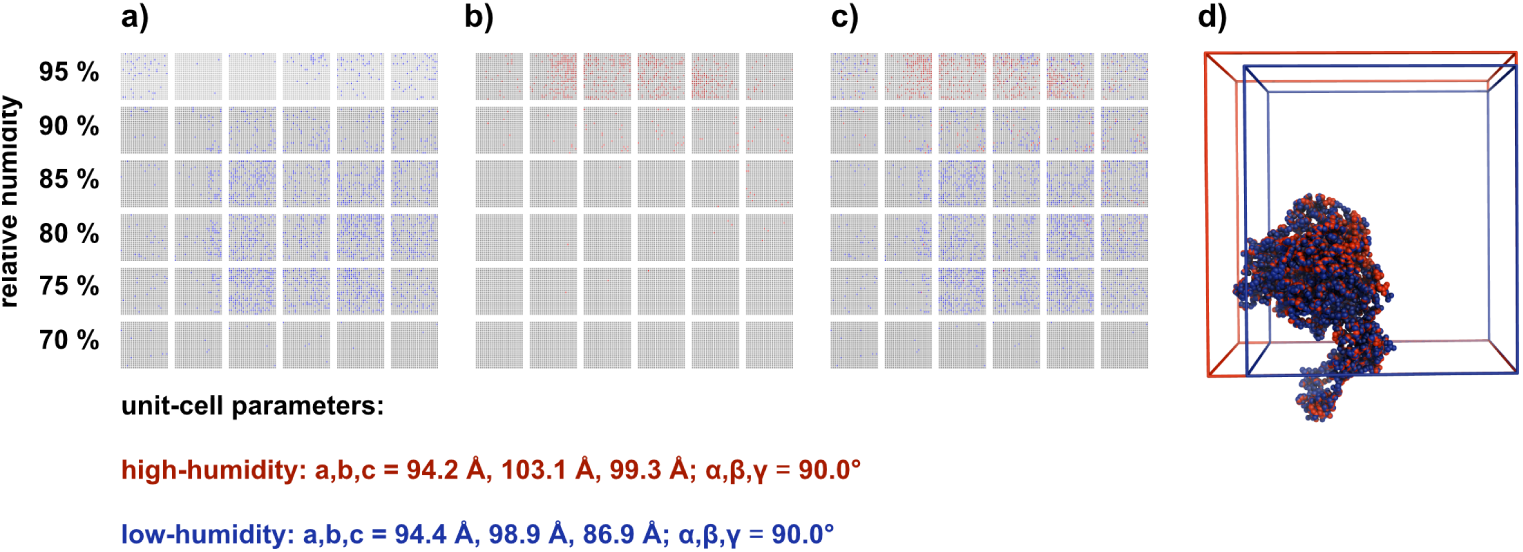
Humidity dependent unit-cell modulation displayed in a hit map. The HARE chips consist of 6×6 compartments, divided into 24×24 features. Each feature is indicated by a small square; a blue square indicates a diffraction pattern in the low humidity unit cell, a red square a diffraction pattern in the larger high-humidity unit cell. Humidity has been reduced by 5% for each row of compartments. a) low-humidity diffraction patterns b) high-humidity diffraction patterns; c) overlay of low- and high-humidity diffraction pattern hits; d) comparison of high- and low-humidity unit-cells.

This emphasizes the sensitivity of protein micro-crystals to environmental humidity, which must be precisely controlled to maintain their diffraction properties. On the other hand, this also opens the opportunity for crystals with large unit cells to be specifically dehydrated to modulate their diffraction properties. The response of protein crystals hydration to their environment has long been known [59] and chemical dehydration devices [60, 67–69] as well as dedicated de-humidification devices have successfully been used for this purpose on single, loop-mounted crystals [54, 57, 62]. With our environmental control box these post-crystallization optimization protocols are now open to serial crystallography.

#### Multi-temperature SSX

To establish that the influence of temperature on the resting state of the CTX-M-14, and XI structures can be accurately determined, initial data were collected without triggering a reaction. After equilibration of the respective temperatures, all parameters were kept equivalent between chips (humidity, beamline, data-collection), and structures were determined using crystals from the same batch of crystals. Five structure of CTX-M-14 were determined at 10 °C, 20 °C, 30 °C, 40 °C, and 50 °C, and five structures of XI at 20°C, 30 °C, 40 °C, 45 °C, and 50 °C (Ext. Tab. 5, 6). This systematic normalization of all experimental parameters, permits direct side-by-side comparison of the structures and consequently allows to directly assign temperature-induced structural differences. To this end we addressed the global structural differences of the individual structures, by fitting the ADPs to a shifted inverse gamma distribution (SIGD) (**Figure 1**). A marked response can be seen in the SIGD of CTX-M-14 once the temperature is increased to 50 °C, which matches the unfolding temperature determined in solution and its pairwise C*_α_RMSD* (see below). By contrast, the SIGD of the XI structures show a gradual response to increasing temperatures. An interesting deviation from the trend can be seen in the 45 °C structure, however, the same pattern can be observed in RoPE-space as well as in the pairwise C*_α_RMSD*, indicating a general structural response in this structure. Taking the general trend into account, the combination with the much higher unfolding temperature of XI in solution, and the gradual increase in ADPs with increasing temperature suggests a different temperature response mechanism of the hyperthermophile XI than the seen in the mesophilic CTX-M-14 protein (see main text).

#### Melting point determination by differential scanning flurometry

To correlate the global structural differences seen in the SIGD with an in-solution behaviour of our model systems, we determined their melting points *via* differential scanning flurometry (Ext. Fig. 6) in four different buffer systems, also taking the crystallization condition into account (see below). The melting point of CTX-M-14 (ca. 53 °C) matches the displacement of the SIGD for the 50 °C SSX structure. By contrast, the multi-temperature SSX experiments for XI do not reach the melting point of XI (ca. 83 °C), but continuously increasing ADPs of XI can be observed in the SIGD.

**Ext. Tab. 4.** Nano-DSF buffer for T*_M_* determination.

|  |  |
| --- | --- |
| CTX-M-14-buffer 1 | 20 mM MES pH 6.0, 50 mM NaCl |
| CTX-M-14-buffer 2 | 140 mM $\text{Li}_2\text{SO}_4$, 15 mM NaCl, 70 mM NaOAc pH 4.5, 6 mM MES, 28% PEG 8000 |
| CTX-M-14-buffer 3 | 140 mM $\text{Li}_2\text{SO}_4$, 15 mM NaCl, 70 mM NaOAc pH 4.5, 6 mM MES |
| CTX-M-14-buffer 4 | 140 mM $\text{Li}_2\text{SO}_4$, 15 mM NaCl, 70 mM NaOAc pH 4.5, 6 mM MES, 1.5% PEG 8000 |
| XI-buffer 1 | 20 mM HEPES pH 7.5 |
| XI-buffer 2 | 10 mM HEPES pH 7.5, 0.2 M $\text{Li}_2\text{SO}_4$, 25% PEG 3350 |
| XI-buffer 3 | 10 mM HEPES pH 7.5, 0.2 M $\text{Li}_2\text{SO}_4$ |
| XI-buffer 4 | 10 mM HEPES pH 7.5, 0.05 M $\text{MgCl}_2$ |

**Ext. Fig. 6.**
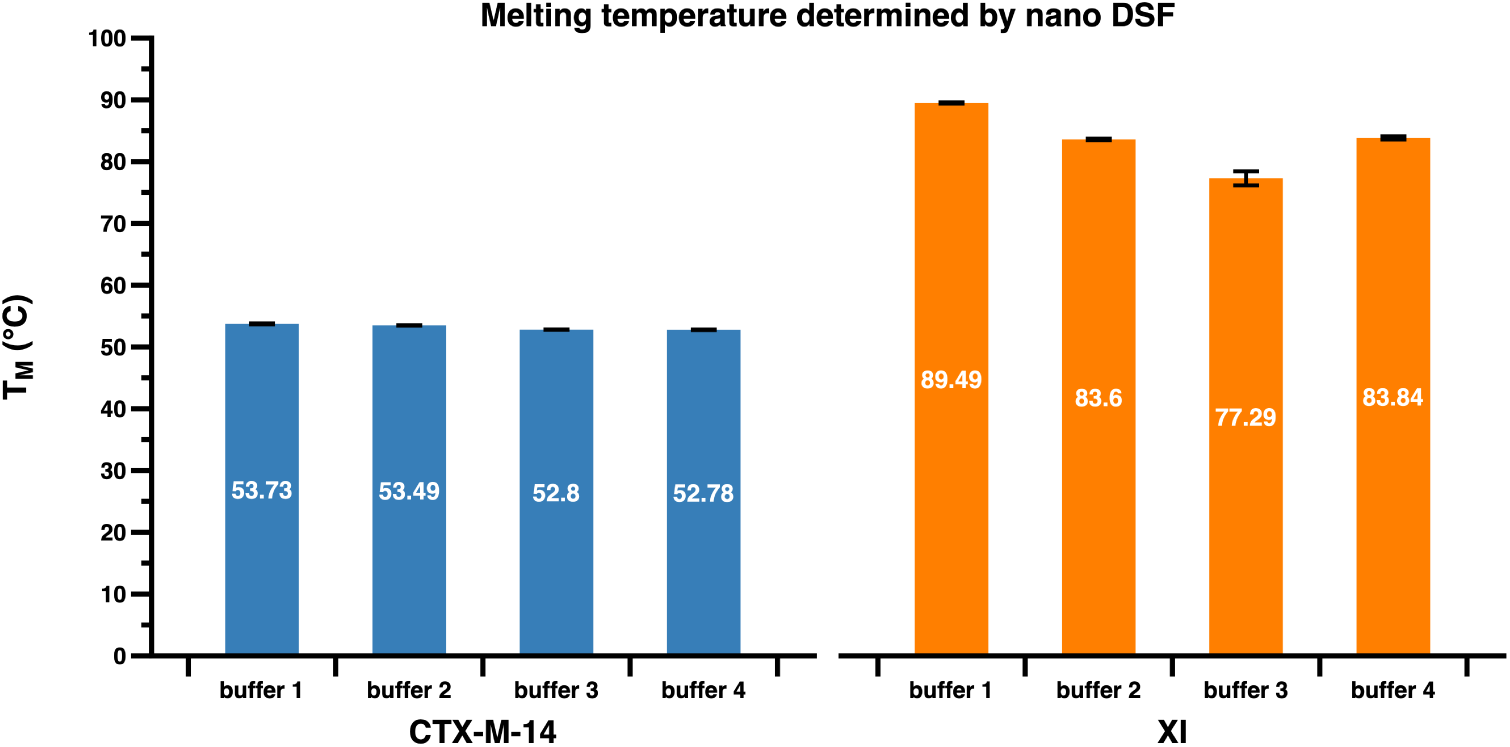
Melting temperature determination. The figure shows the melting temperature for CTX-M-14 and XI determined by nanoDSF in 4 different buffer systems each. The in-solution melting temperature reflects the activity optima of the mesophilic and hyperthermophilic proteins, respectively.

#### C*_α_* RMSD

In order to quantify and visualise the changing global structural differences, a pairwise backbone Root Mean Square Deviation (C*α* RMSD) was calculated for both model systems and represented as a categorical heatmap (Ext.Fig. 7). Before calculating the RMSD for a pair, the two structures were aligned *via* Singular Value Decomposition (SVD) using the Biopython package Bio.SVDSuperimposer in order to minimise the resulting RMSD value. A temperature dependent clustering is observed for both model systems.

**Ext. Fig. 7.**
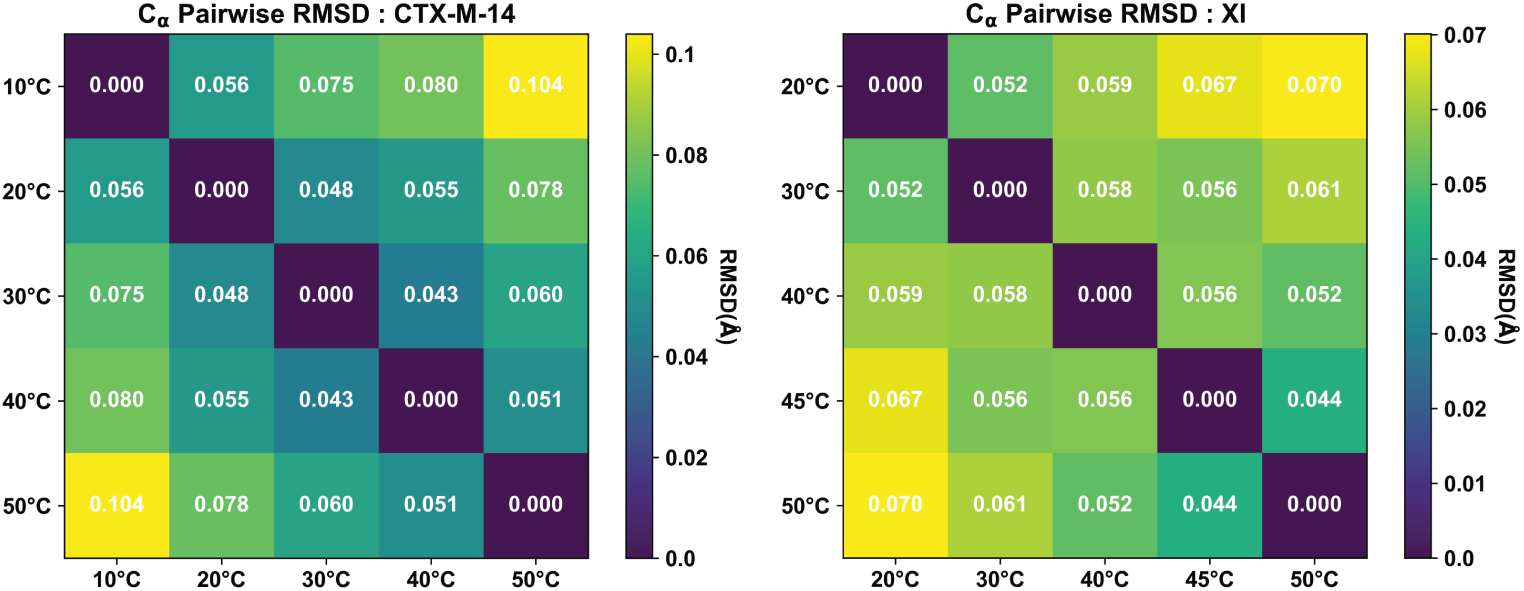
Pairwise C*α* RMSD heatmap. Backbone RMSDs in Å between pairs of structures at different temperatures for both model systems.

#### Multi-temperature time-resolved SSX

To study temperature effects on catalytic turnover we determined three CTX-M-14 crystal structures 3 s after reaction initiation with piperacillin at 20 °C, 30 °C and 37 °C, respectively (Ext. Tab. 7), and for XI we determined five crystal structures 60 s after reaction initiation with glucose at 20 °C, 30 °C, 40 °C, 45 °C and 50 °C, respectively (Extended data table 8). Any time-resolved crystallographic analysis will result in a superimposed mixture of sub-states, generating an ensemble of structures with respective fractional occupancies [25]. To address this situation and minimize interpretation bias, we initially assembled all previously known stable intermediates into one structure and relied on constrained group refinement to determine the occupancy of the different overlapping states at the respective datasets [26]. For CTX-M-14 the 20 °C structure corresponds to the unbound state, exhibited by rather discontinuous difference electron density and accordingly low occupancy for any of the piperacillin intermediates. The 30 °C structure on the other hand is clearly populated by ligand difference density. Occupancy refinement indicated a mixture that predominantly corresponds to the acyl-enzyme intermediate and a complex with the hydrolysed piperacillin product. Finally, the 37 °C structure mainly corresponds to CTX-M-14 in complex with the hydrolysed piperacillin product (**Fig. 2 a-c**).

**Ext. Tab. 5.**
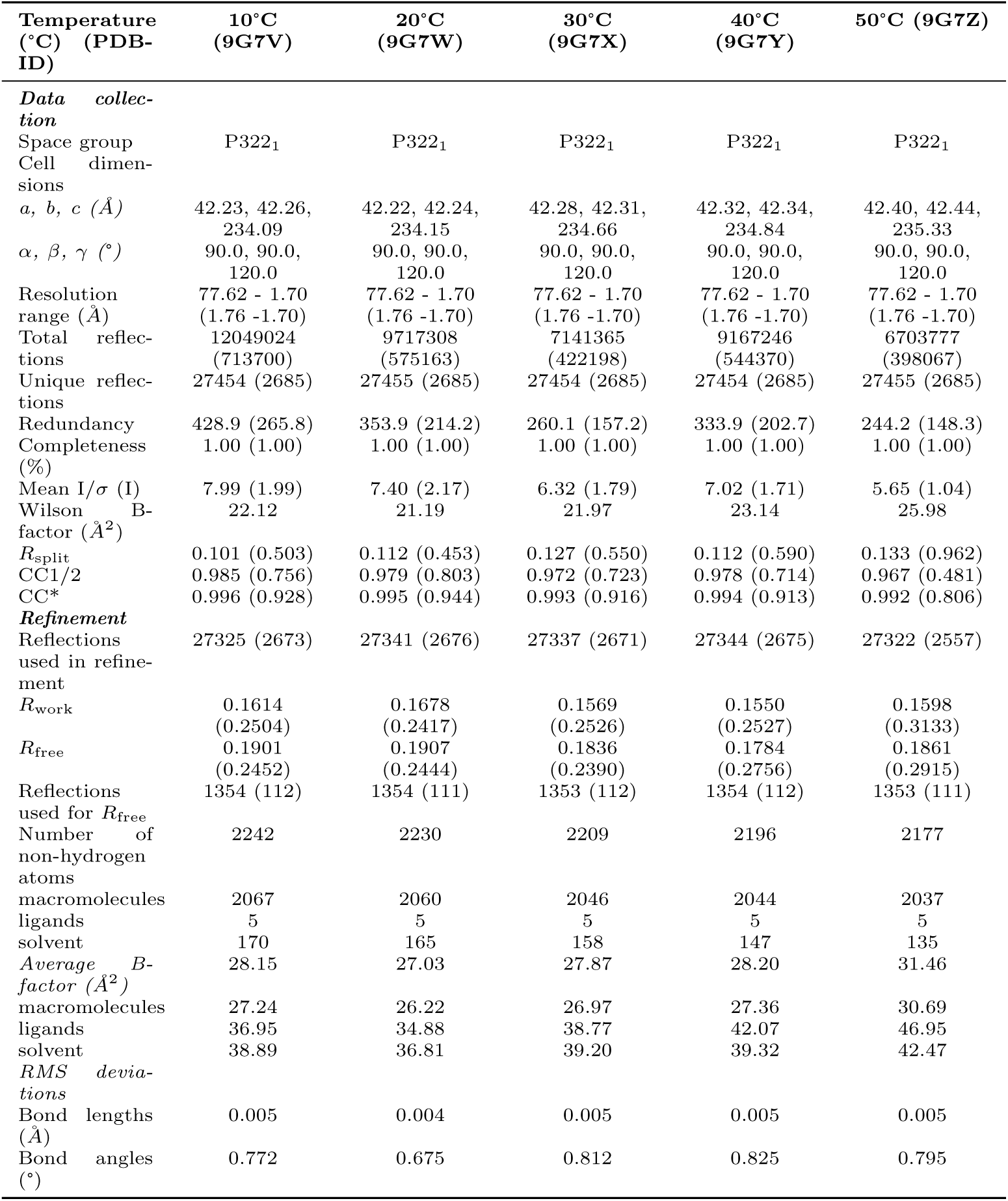
Data collection and refinement statistics for CTX-M-14 apo data. Values in the highest resolution shell are shown in parentheses.

For XI on the other hand, we modelled the unbound state, the Michaelis-Menten complex with closed-ring glucose as well as an open-chain glucose intermediate consistent with previously determined structures. The respective substrate complex structure at 20°C can be superimposed with an RMSD. of 0.1 Å to previously reported substrate complexes at room temperature (PDB-ID: 3KCL) [70] (Fig. 2). The multiple overlapping states make an unambiguous interpretation based on the electron density alone difficult, which mandates the use of more complex modelling and analysis protocols in future experiments [26, 71, 72]. However, by increasing the temperature, different catalytic states (*i.e.*, closed and open-chain intermediates) can be captured at the same delay time, clearly indicating an increase in the catalytic activity of XI with increasing temperature.

**Ext. Tab. 6.** Data collection and refinement statistics of the XI apo data. *Values in the highest resolution shell are shown in parentheses*.

| Temperature<br>(°C) (PDB-ID) | 20°C<br>(9G5N) | 30°C (9G5S) | 40°C<br>(9G5W) | 45°C<br>(9G5X) | 50°C (9G61) |
| --- | --- | --- | --- | --- | --- |
| *Data collection* |  |  |  |  |  |
| Space group | I 2 2 2 | I 2 2 2 | I 2 2 2 | I 2 2 2 | I 2 2 2 |
| Cell dimensions |  |  |  |  |  |
| *a*, *b*, *c* (Å) | 94.2 103.05<br>99.25 | 94.2 103.05<br>99.25 | 94.2 103.05<br>99.25 | 94.2 103.05<br>99.25 | 94.2 103.05<br>99.25 |
| $\alpha$, $\beta$, $\gamma$ (°) | 90.0 90.0 90.0 | 90.0 90.0 90.0 | 90.0 90.0 90.0 | 90.0 90.0 90.0 | 90.0 90.0 90.0 |
| Resolution | 71.49 - 1.7 | 71.49 - 1.7 | 71.49 - 1.7 | 71.49 - 1.7 | 71.49 - 1.7 |
| range (Å) | (1.761 - 1.7) | (1.761 - 1.7) | (1.761 - 1.7) | (1.761 - 1.7) | (1.761 - 1.7) |
| Total reflections | 20298653 | 10229855 | 10613133 | 81189855 | 23289164 |
| Unique reflections | 53351 (5284) | 53338 (5277) | 53317 (5271) | 53331 (5273) | 53297 (5253) |
| Redundancy | 380.3(224.3) | 191.7(113.2) | 198.8(117.7) | 1521.2(904.5) | 436.3(258.8) |
| Completeness (%) | 99.96 (99.96) | 99.93 (99.83) | 99.89 (99.72) | 99.92 (99.75) | 99.85 (99.38) |
| Mean I/$\sigma$(I) | 5.55(3.31) | 3.83(1.50) | 3.59(0.98) | 8.57(2.88) | 4.87(1.14) |
| Wilson B-factor | 15.26 | 17.54 | 20.14 | 19.82 | 21.96 |
| $R_{split}$ | 20.92(33.98) | 26.71(66.18) | 24.74(102.55) | 10.09(31.25) | 17.46(90.39) |
| $CC^{1/2}$ | 92.76(76.23) | 89.39(56.02) | 92.27(38.03) | 98.64(86.29) | 96.25(45.20) |
| $CC^*$ | 98.10(93.01) | 97.16(84.74) | 97.97(74.23) | 99.66(96.25) | 99.04(78.90) |
| *Refinement* |  |  |  |  |  |
| Reflections used in refinement | 53351 (5284) | 53338 (5277) | 53317 (5271) | 53331 (5273) | 53297 (5253) |
| $R_{work}$ | 0.1624<br>(0.2901) | 0.1680<br>(0.3272) | 0.1727<br>(0.3735) | 0.1453<br>(0.3158) | 0.1590<br>(0.3597) |
| $R_{free}$ | 0.1906<br>(0.3290) | 0.2096<br>(0.3910) | 0.2072<br>(0.4166) | 0.1824<br>(0.3611) | 0.1903<br>(0.3987) |
| Reflections used for $R_{free}$ | 2607 (291) | 2607 (291) | 2606 (291) | 2605 (290) | 2603 (288) |
| Number of non-hydrogen atoms | 3637 | 3566 | 3638 | 3552 | 3543 |
| macromolecules | 3275 | 3234 | 3325 | 3261 | 3284 |
| ligands | 2 | 3 | 2 | 3 | 2 |
| solvent | 360 | 329 | 311 | 288 | 257 |
| Average B-factor (Å<sup>2</sup>) | 17.69 | 20.2 | 23.47 | 22.92 | 25.36 |
| macromolecules | 16.22 | 19.02 | 22.33 | 21.86 | 24.56 |
| ligands | 16.02 | 11.48 | 17.73 | 12.44 | 18.2 |
| solvent | 31.14 | 31.91 | 35.64 | 35.04 | 35.67 |
| *RMS deviations* |  |  |  |  |  |
| Bond lengths (Å) | 0.006 | 0.016 | 0.004 | 0.013 | 0.004 |
| Bond angles (°) | 0.8 | 1.34 | 0.65 | 1.23 | 0.67 |

**Ext. Tab. 7.** Data collection and refinement statistics of the CTX-M-14 Piperacillin data at a time-delay of 3s. Values in the highest resolution shell are shown in parentheses.

| Temperature (°C) (PDB-ID) | 20°C (9G80) | 30°C (9G81) | 37°C (9G82) |
| --- | --- | --- | --- |
| **Data collection** |  |  |  |
| Space group | P322<sub>1</sub> | P322<sub>1</sub> | P322<sub>1</sub> |
| Cell dimensions |  |  |  |
| *a*, *b*, *c* (Å) | 42.15, 42.15,<br>234.14 | 42.20, 42.20,<br>234.53 | 42.25, 42.25,<br>234.85 |
| $\alpha, \beta, \gamma$ (°) | 89.99, 89.99,<br>119.98 | 89.99, 89.99,<br>119.93 | 89.99, 89.98,<br>119.93 |
| Resolution range (Å) | 78.07 - 1.70 (1.76<br>-1.70) | 78.07 - 1.70 (1.76<br>-1.70) | 78.07 - 1.70 (1.76<br>-1.70) |
| Total reflections | 1266008 (75053) | 3599661 (216133) | 4883576 (290896) |
| Unique reflections | 27981 (2739) | 27439 (2662) | 27982 (2734) |
| Redundancy | 45.2 (27.4) | 131.2 (81.2) | 174.5 (106.4) |
| Completeness (%) | 1.00 (1.00) | 1.00 (1.00) | 1.00 (1.00) |
| Mean I/$\sigma$ (I) | 1.91 (0.25) | 4.46 (1.04) | 4.36 (0.66) |
| Wilson B-factor (Å<sup>2</sup>) | 26.44 | 22.25 | 24.78 |
| R<sub>split</sub> | 0.364 (4.577) | 0.182 (0.995) | 0.163 (1.595) |
| CC1/2 | 0.862 (0.034) | 0.952 (0.446) | 0.959 (0.252) |
| CC* | 0.962 (0.257) | 0.988 (0.786) | 0.990 (0.634) |
| **Refinement** |  |  |  |
| Reflections used in refinement | 26826 (1948) | 27336 (2363) | 27851 (2569) |
| R<sub>work</sub> | 0.2281 (0.4130) | 0.1645 (0.3038) | 0.1687 (0.3592) |
| R<sub>free</sub> | 0.2556 (0.4706) | 0.1948 (0.3061) | 0.1995 (0.3746) |
| Reflections used for R<sub>free</sub> | 1315 (103) | 1354 (141) | 1377 (154) |
| Number of non-hydrogen atoms | 2362 | 2371 | 2408 |
| macromolecules | 2156 | 2148 | 2193 |
| ligands | 41 | 41 | 41 |
| solvent | 165 | 182 | 174 |
| Average B-factor (Å<sup>2</sup>) | 36.58 | 29.28 | 33.14 |
| macromolecules | 35.32 | 28.21 | 31.89 |
| ligands | 49.55 | 30.60 | 39.74 |
| solvent | 43.45 | 40.71 | 44.02 |
| **RMS deviations** |  |  |  |
| Bond lengths (Å) | 0.002 | 0.005 | 0.003 |
| Bond angles (°) | 0.861 | 1.046 | 0.949 |

### Code

#### Script for the Shifted inverse gamma distribution

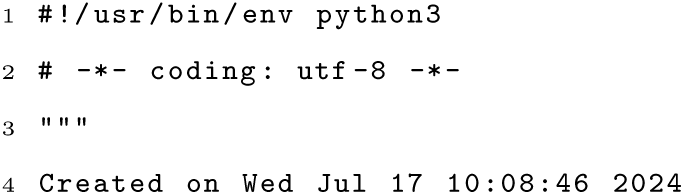
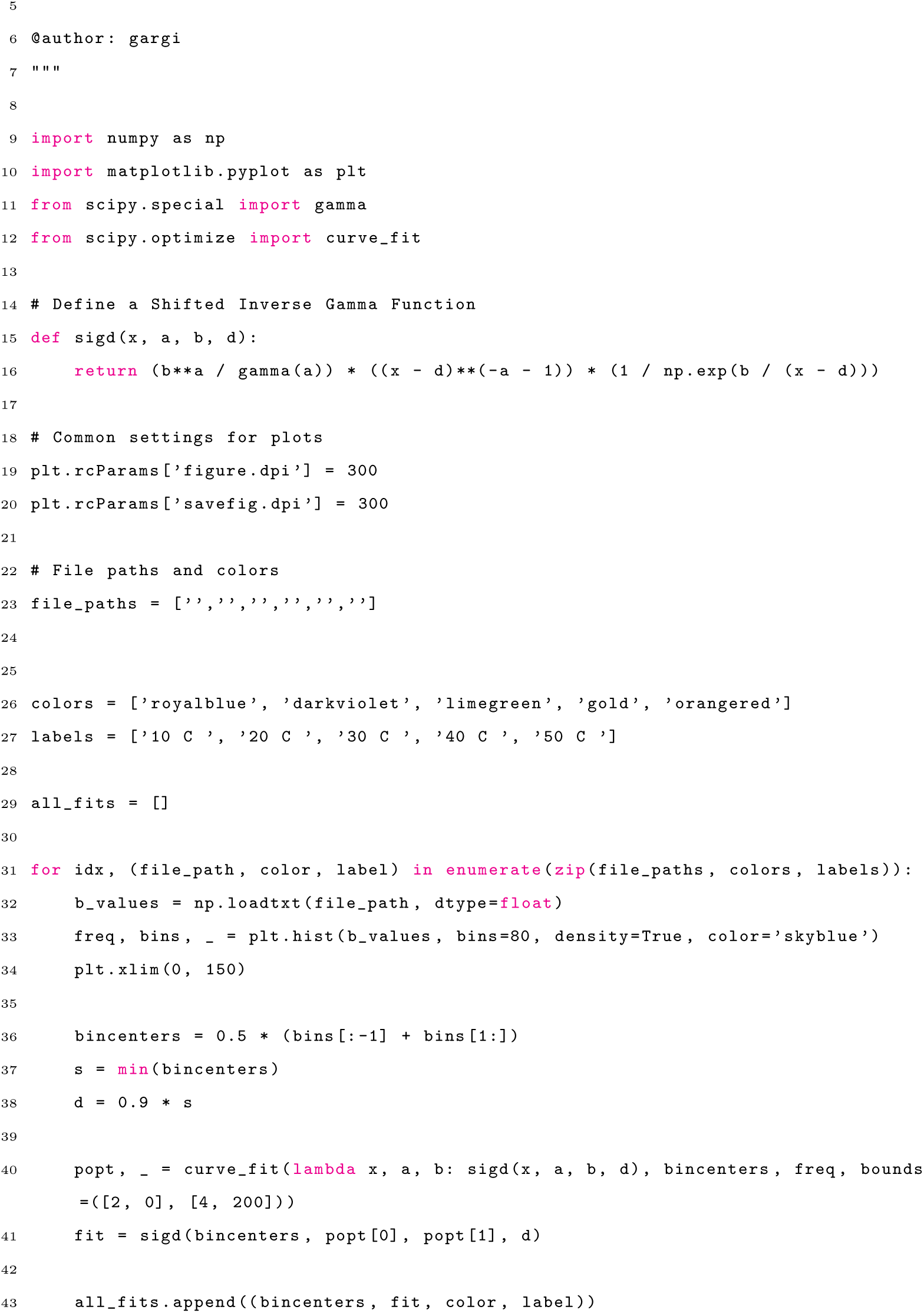
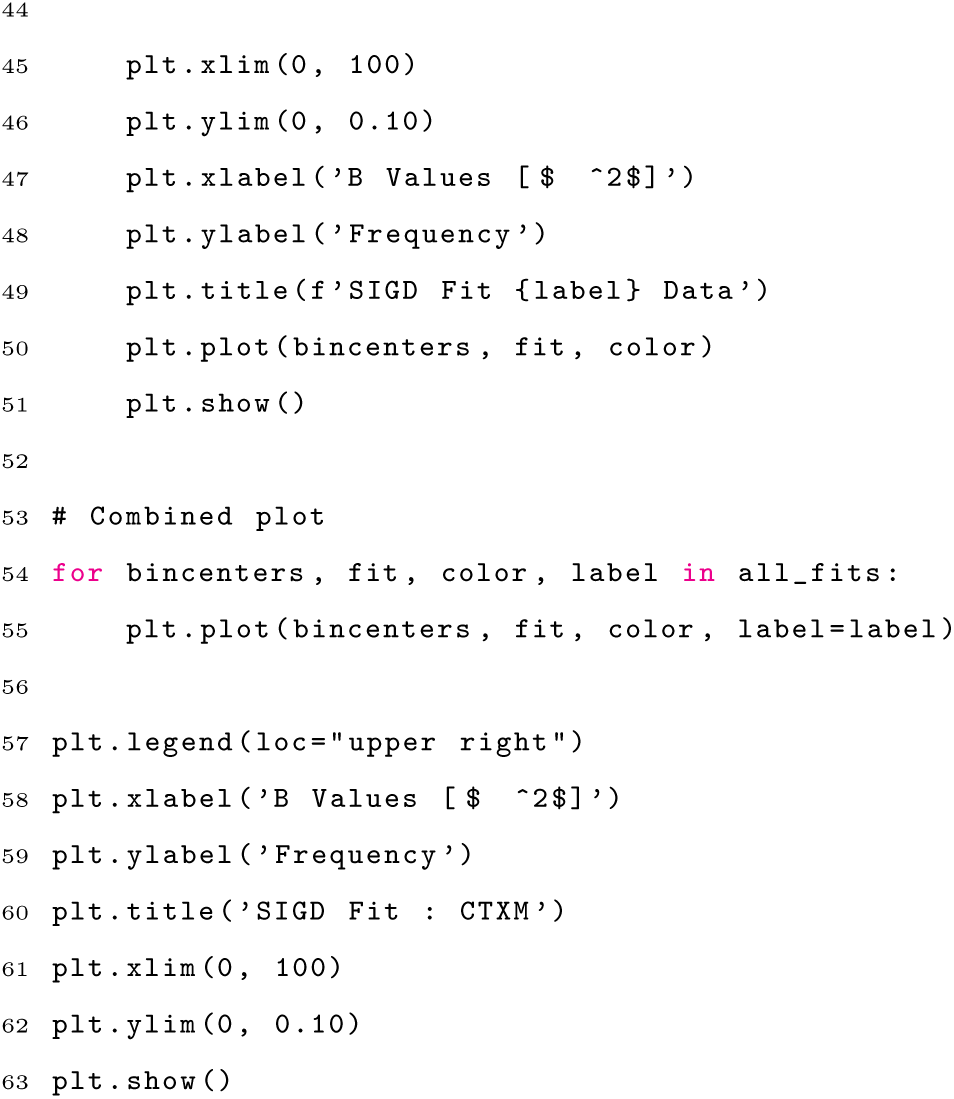

**Ext. Tab. 8.** Data collection and refinement statistics of the XI 60s data. *Values in the highest resolution shell are shown in parentheses*.

| Temperatutre<br>(°C) (PDB-ID) | 20°C<br>(9G6L) | 30°C<br>(9G6M) | 40°C<br>(9G6N) | 45°C<br>(9G6O) | 50°C<br>(9G6P) |
| --- | --- | --- | --- | --- | --- |
| *Data collec-*<br>*tion* |  |  |  |  |  |
| Space group | I 2 2 2 | I 2 2 2 | I 2 2 2 | I 2 2 2 | I 2 2 2 |
| Cell dimen-<br>sions |  |  |  |  |  |
| *a, b, c* (Å) | 94.2 103.05<br>99.25 | 94.2 103.05<br>99.25 | 94.2 103.05<br>99.25 | 94.2 103.05<br>99.25 | 94.2 103.05<br>99.25 |
| $\alpha, \beta, \gamma$ (°) | 90.0 90.0 90.0 | 90.0 90.0 90.0 | 90.0 90.0 90.0 | 90.0 90.0 90.0 | 90.0 90.0 90.0 |
| Resolution | 71.49 - 1.7 | 71.49 - 1.7 | 71.49 - 1.7 | 71.49 - 1.7 | 71.49 - 1.7 |
| range (Å) | (1.761 - 1.7) | (1.761 - 1.7) | (1.761 - 1.7) | (1.761 - 1.7) | (1.761 - 1.7) |
| Total reflec-<br>tions | 9415749 | 12055922 | 11970298 | 26069844 | 8506453 |
| Unique<br>reflections | 53357 (5281) | 53350 (5281) | 53330 (5270) | 53342 (5278) | 53317 (5273) |
| Redundancy | 176.4(104.5) | 225.9(134.2) | 224.3(133.2) | 488.4(290.4) | 159.4(94.7) |
| Completeness<br>(%) | 99.97 (99.91) | 99.95 (99.91) | 99.92 (99.70) | 99.94 (99.85) | 99.89 (99.75) |
| Mean I/$\sigma$(I) | 4.37(2.68) | 4.99(2.46) | 4.42(1.65) | 6.39(2.67) | 3.24(0.93) |
| Wilson B-factor | 15.89 | 17.92 | 19.59 | 21.24 | 22.4 |
| CC<sup>1/2</sup> | 89.59(73.39) | 92.56(76.39) | 93.27(62.3) | 96.44(79.42) | 89.5(36.2) |
| CC* | 97.22(92.01) | 98.05(93.07) | 98.24(87.62) | 99.09(94.09) | 97.19(72.91) |
| *Refinement* |  |  |  |  |  |
| Reflections<br>used in<br>refinement | 53357 (5281) | 53350 (5281) | 53330 (5270) | 53342 (5278) | 53317 (5273) |
| Reflections<br>used for R-free | 2608 (291) | 2607 (290) | 2605 (289) | 2608 (291) | 2606 (290) |
| *work* | 0.1657<br>(0.2765) | 0.1549<br>(0.2899) | 0.1604<br>(0.3401) | 0.1479<br>(0.2998) | 0.1744<br>(0.3566) |
| *free* | 0.2051<br>(0.3178) | 0.1922<br>(0.3259) | 0.1997<br>(0.4004) | 0.1859<br>(0.3468) | 0.2153<br>(0.3938) |
| *Number of*<br>*non-hydrogen*<br>*atoms* | 3617 | 3553 | 3590 | 3568 | 3507 |
| macromolecules | 3259 | 3224 | 3282 | 3270 | 3221 |
| ligands | 27 | 27 | 27 | 27 | 27 |
| solvent | 331 | 302 | 281 | 271 | 259 |
| *Average B-*<br>*factor (Å<sup>2</sup>)* | 19.03 | 21 | 23.39 | 24.18 | 26.14 |
| macromolecules | 17.77 | 19.93 | 22.44 | 23.3 | 25.37 |
| ligands | 14.24 | 16.09 | 18.23 | 19.95 | 21.63 |
| solvent | 31.89 | 32.89 | 35.02 | 35.25 | 36.23 |
| *RMS devia-*<br>*tions* |  |  |  |  |  |
| Bond lengths<br>(Å) | 0.011 | 0.014 | 0.006 | 0.017 | 0.006 |
| Bond angles<br>(°) | 1.09 | 1.29 | 0.8 | 1.49 | 0.78 |

#### Script to calculate the C*_α_* RMSD

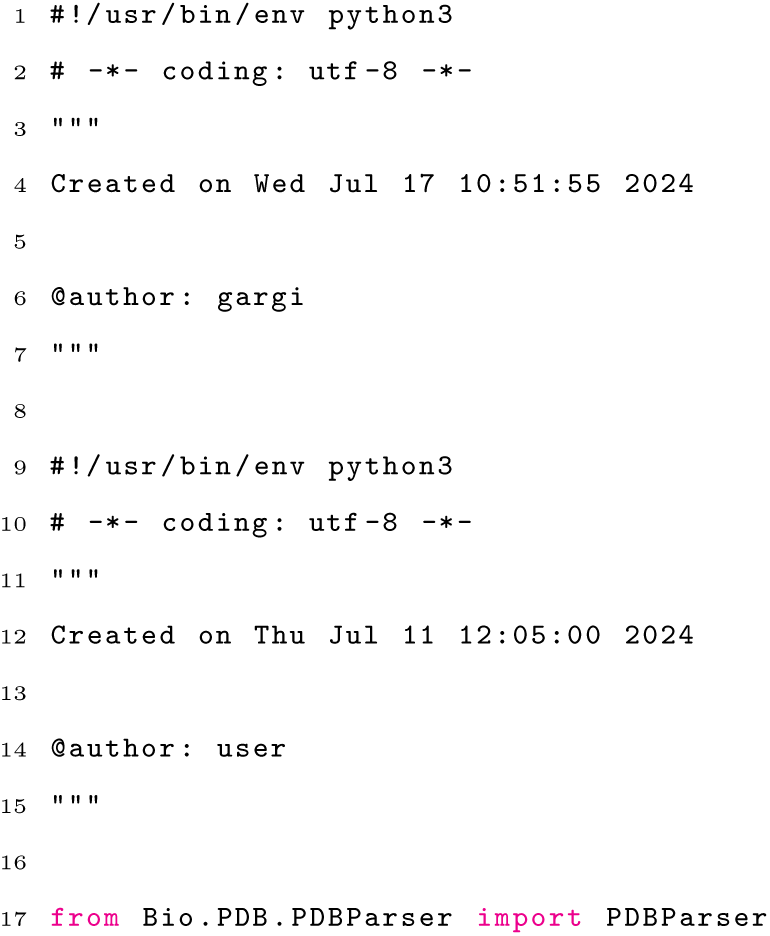
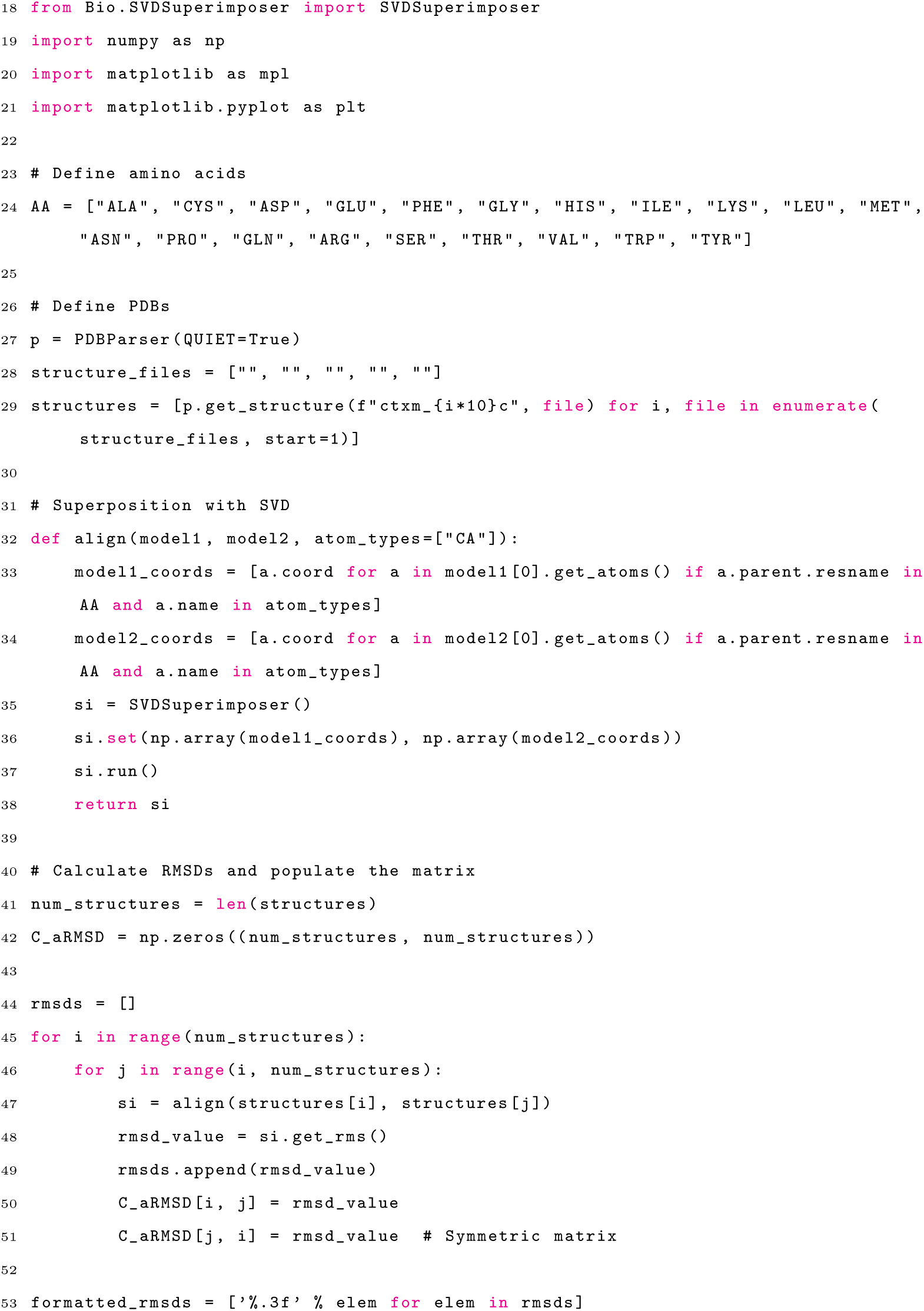
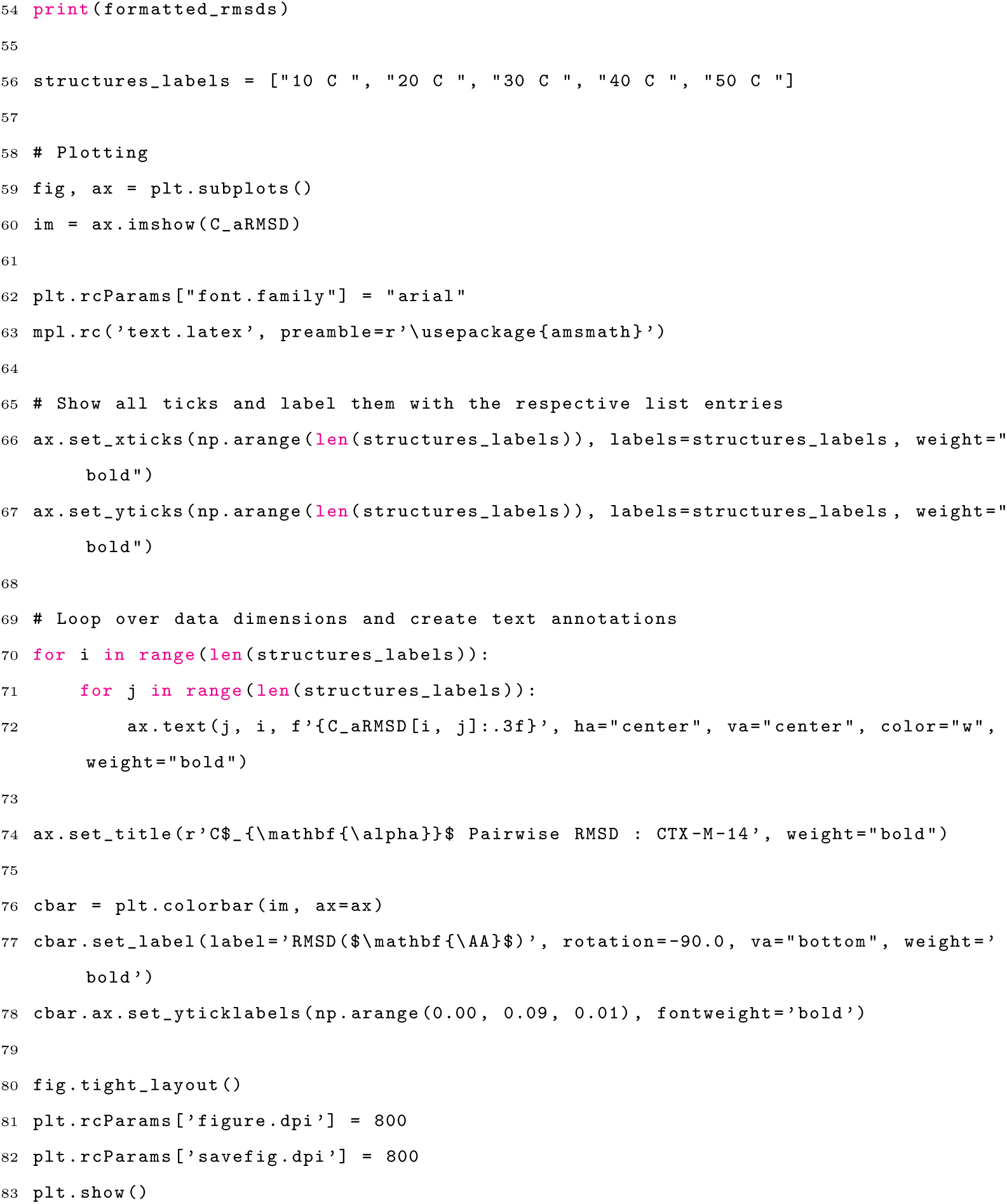

